# Input-specific bi-directional regulation of CA3 pyramidal cell excitability: its implications in sequence learning

**DOI:** 10.1101/2023.03.29.534692

**Authors:** Kisang Eom, Yujin Kim, Seungmin Baek, Alan J Park, Won-Kyung Ho, Suk-Ho Lee

**Affiliations:** Department of Physiology, Seoul National University College of Medicine, 103 Daehak-ro, Jongno-gu, 03080, Seoul, Republic of Korea; Department of Brain and Cognitive Science, Seoul National University College of Natural Science, 103 Daehak-ro, Jongno-gu, 03080, Seoul, Republic of Korea; Department of Physiology, Keimyung University School of Medicine, 1095 Dalgubeol-daero street, Dalseo-gu district, Daegu, Republic of Korea

**Author notes:** Correspondence: Suk-Ho Lee.

**Keywords:** CA3 pyramidal cell, intrinsic excitability, depotentiation, neuronal ensemble, computational modelling, sequence learning

## Abstract

Neuronal excitability is a key determinant for recruitment of a neuron to an ensemble. High-frequency mossy fiber (MF) inputs induce a prolonged increase in the excitability of a CA3 pyramidal cell (called long-term potentiation of intrinsic excitability, LTP-IE), thereby weak perforant pathway (PP) inputs can induce long-term potentiation at PP synapses (PP-LTP). However, sustained hyperexcitability is detrimental, and a mechanism to reverse this primed state is necessary. Here, we show that burst firings of CA3 pyramidal cells elicited by PP or recurrent synaptic inputs reverse the MF-induced LTP-IE. Moreover, the high-frequency PP inputs to MF-primed CA3 pyramidal cells induced not only PP-LTP but also restored the high excitability state. Labeling a neuronal ensemble using c-fos promoter in animals exposed to a novel context, we found most CA3 ensemble cells exhibited increased excitability, indicative of LTP-IE. Moreover, when the animals experienced novel contexts twice with an interval, a substantial subset of putative twice-activated CA3 ensemble cells exhibited reduced excitability, implying depotentiation of LTP-IE. We developed an in silico model based on these experimental results and found that MF-induced LTP-IE and its depotentiation are critical for association of orthogonal neuronal ensembles representing temporally discontiguous events.

**Highlights:**

- It is unknown how non-overlapping ensembles are linked in the hippocampal CA3 area.
- Mossy fiber inputs prime CA3 pyramidal cells by enhancing dendritic excitability.
- Perforant pathway (PP) inputs to the primed cells induce synaptic strengthening.
- At the same time, the high excitability state is restored by PP inputs.
- This learning rule may play a key role in sequence learning in CA3 network.

## Introduction

Hippocampal CA3 pyramidal cells (CA3-PCs) receive two kinds of extrinsic synaptic inputs from dentate gyrus and entorhinal cortex via mossy fibers (MFs) and perforant pathway (PP), respectively. Sparse and strong non-Hebbian MF inputs contribute to orthogonal encoding, while weak and a larger number of Hebbian PP inputs are required for cued memory retrieval of a pattern stored in the CA3 recurrent network (Treves and Rolls 1992; O’Reilly and McClelland, 1994; Lee & Kesner, 2004). Involvement of PP synapses in retrieval of episodic memory implies that long-term plasticity (LTP) at PP synapses would play a key role in triggering reactivation of CA3 ensembles from auto-associative CA3 network during cued retrieval. Because PP synaptic inputs are not strong enough to drive firing of CA3-PCs, MF-driven firing would be required for induction of Hebbian LTP at PP synapses (McNaughton and Morris, 1987). High frequency MF inputs not only provide such immediate depolarization but also induce sustained enhancement of dendritic voltage responses to PP synaptic inputs, called long-term potentiation of intrinsic excitability (LTP-IE) (Eom et al., 2019; Hyun et al., 2015; Hyun et al., 2013). The MF-induced LTP-IE is mediated by an increase in the distal apical dendritic excitability (Hyun et al., 2013), and potentiates the EPSP-to-spike (E-S) coupling of PP inputs (E-S potentiation) (Hyun et al., 2015). Therefore, high-frequency MF inputs prime a CA-PC for homosynaptic LTP at PP synapses (PP-LTP) because the E-S potentiation lowers the threshold of PP input strength required for induction of PP-LTP (called LTP threshold) (Eom et al. 2022). Given that LTP-IE is not a synapse-specific process but occurs over a whole distal dendritic arbor (Hyun et al., 2013), LTP-IE without depotentiation would lead to rapid saturation of PP-CA3 synaptic weights, which will impair further learning (Moser et al., 1998), and excitation/inhibition imbalance of the CA3 recurrent network. It remains unexplored, however, how MF-induced high excitability state of CA3-PC is restored to the baseline.

The other question to be addressed is whether LTP-IE indeed occur in CA3-PCs that participate in neuronal ensembles (called CA3 ensemble cells) *in vivo*. This question is crucial to understand CA3 ensemble dynamics because neuronal excitability is a key factor that determines whether a given neuron is recruited to an ensemble carrying information. Once a CA3-PC receives high-frequency MF inputs and undergoes LTP-IE in a previous episode, the probability for the primed cell to be recruited to an ensemble in the next episode would be enhanced because E-S potentiation would enable PP input alone to drive firing in the primed CA3-PC. Whereas dynamic changes of intrinsic excitability accompanied with behavioral learning have been explored in hippocampal DG and CA1 area (Oh & Disterhoft, 2015; Pignatelli et al., 2019),the same issue in the CA3 region remains unexplored. The present study was set out to address these two questions. Here, we show that MF input-induced primed state of a CA3-PC is terminated by high-frequency PP or associational/commissural (A/C) inputs. Moreover, CA3 ensemble cells exhibited indeed high excitability features similar to CA3-PCs that underwent LTP-IE, and LTP-IE was reversed in a subset of putative twice-activated CA3 ensemble cells when an animal experienced two temporally separated events. Incorporating in vitro findings to the Hebb-Marr CA3 network model (McNaughton & Morris, 1987), we propose that the MF- and PP-induced bi- directional regulation of CA3-PC excitability may link two orthogonal ensembles representing two events adjacent in a sequence even if they are temporally separated.

## Materials and Methods

### Animals and ethical approval

All studies, experimental protocols, and animal manipulation protocol described in this article were conducted with the Institutional Animal Care and Use Committee (IACUC). The animals were maintained in standard environmental conditions (Temperature: 25 ± 2 °C, Humidity: 60 ± 5 %, Dark/Light cycle: 12/12h) and monitored under veterinary supervision by Institute for Experimental Animals, Seoul National University College of Medicine.

### Preparation of hippocampal slices

Acute transverse hippocampal slices were obtained from Sprague-Dawley rats (PW2-3 for Figures 1-4 and Figures S2-S3; PW6 for Figure 5; PW, postnatal weeks) or cfos-shEGFP mice (PW6-8 for Figures 6-7) or ZnT-heteroKO and ZnT-heteroKO/cfos-shEGFP mice (PW6, Figure 8) of either sex. Animals were anesthetized by inhalation of isoflurane. After decapitation, brain was quickly removed and chilled in ice-cold preparation solution containing 75 sucrose, 87 NaCl, 25 NaHCO_3_, 2.5 KCl, 1.25 NaH_2_PO_4_, 25 D-glucose, 7 MgCl_2_, and 0.5 CaCl_2_ (in mM; equilibrated with carbogen mixture of 95% O_2_ and 5% CO_2_). After mounting on a slicer (Leica VT1200, Nussloch, Germany), 300 μm (whole-cell recording)-thick transverse slices (whole-cell recording) were prepared and incubated at 34 °C for 30 min in the preparation solution, and thereafter stored at room temperature (22 °C). For experiments, slices were transferred to a submersion recording chamber superfused with standard aCSF containing 124 NaCl, 26 NaHCO_3_, 3.2 KCl, 1.25 NaH_2_PO_4_, 10 D-glucose, 2.5 CaCl_2_, and 1.3 MgCl_2_ (in mM).

**Fig. 1.**
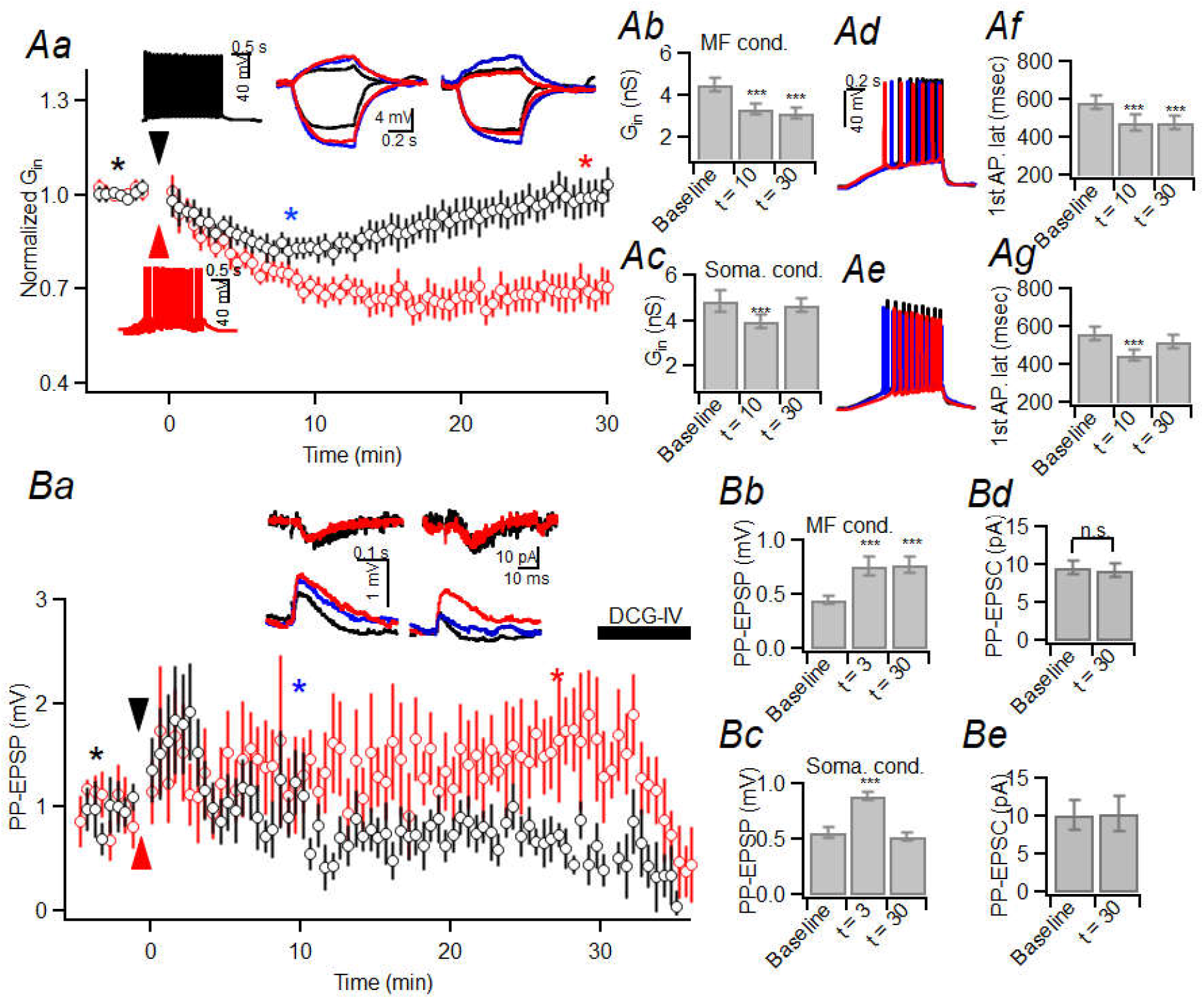
Differential effects of intracellular GSH on somatic and MF conditioning-induced LTP-IE. **Aa** and **Ba.** Mean baseline-normalized input conductance (G_in_) and PP-EPSP amplitude of CA3-PCs before and after MF (red) or somatic (black) conditioning (arrowheads). Right two insets of Aa, Voltage responses to subthreshold current injection for measuring G_in_ before and after MF (middle) and somatic (right) conditioning. Insets of Ba, PP-EPSCs (upper) and PP-EPSPs (lower) before and after MF (left) or somatic (right) conditioning. The time points of recording are indicated by asterisks (same color codes). **Ab-Ac.** Summary for G_in_ at baseline, 10 min and 30 min after the MF (*Ab*) or somatic (*Ac*) stimulation. For MF conditioning, 0 vs. 10 min, p < 0.001; 0 vs. 30 min, p < 0.001; 10 vs. 30 min, p = 0.190 (GLM and simple effect analysis). For somatic conditioning, 0 vs. 10 min, p < 0.001; 0 vs 30 min, p = 0.533; 10 vs. 30 min: p < 0.001 (GLM and simple effect analysis). Here, the baseline values are denoted as 0. **Ad-Ae.** Voltage responses to ramp current injection measured before (black), 10 min (blue) and 30 min (red) after MF (*Ad*) and somatic (*Ae*) conditioning. **Af-Ag.** Mean latency of first AP before, and 10 min and 30 min after the MF (*Af*) or somatic (*Ag*) conditioning (For MF, 0 vs. 10 min, p < 0.001; 0 vs. 30 min, p < 0.001; 10 vs. 30 min, p = 0.964; For somatic, 0 vs. 10 min, p < 0.001; 0 vs. 30 min, p = 0.054; 10 vs. 30 min, p < 0.001; GLM and simple effect analysis). **Bb-Be.** Mean PP-EPSP (Bb-Bc) and PP-EPSC (Bd-Be) amplitudes at baseline, 3 min and 30 min after MF (Bb, Bd) or somatic (Bc, Be) conditioning. For EPSP/MF conditioning, 0 vs. 10 min, p < 0.001; 0 vs. 30 min, p < 0.001; 10 vs. 30 min: p = 0.779. For EPSP/somatic conditioning, 0 vs. 10 min, p < 0.001, 0 vs. 30 min, p = 0.481; 10 vs. 30 min, p < 0.001 (GLM and simple effect analysis).

### Electrophysiological study

Whole-cell voltage- or current-clamp recordings from CA3-PCs were performed at near-physiological temperature (34 ± 1 °C), while the recording chamber was perfused with the standard aCSF at 1 ∼ 1.5 ml/min. Patch pipettes were pulled from borosilicate glass tubing (outer diameter: 1.5 mm, wall thickness: 0.225 mm) with a horizontal pipette puller (P-97, Sutter Instruments, Novato, CA, USA) and filled with the intracellular solution containing 130 K-gluconate, 7 KCl, 1 MgCl_2_, 2 Mg-ATP, 0.3 Na-GTP, 10 HEPES, 0.1 EGTA (in mM; pH adjusted to 7.20 with KOH, 295 mOsm) with the pipette resistance was 3 ∼ 4 MΩ. After formation of whole-cell configuration on the somata of CA3-PCs, recordings were performed from only those cells that had stable resting membrane potential within −76 and −58 mV (cells that had more positive resting membrane potential or unstable resting membrane potential were discarded). Under this condition, input conductance (G_in_) was measured from subthreshold voltage responses to −30 pA and +10 pA current steps of 0.5 s (Hyun et al., 2015). G_in_ was monitored every 10 s before and after delivery of a conditioning protocol. Recordings were obtained in the presence of a GABA_A_ receptor antagonist, picrotoxin (PTX, 100 μM) unless specified. All recordings were made using a MultiClamp 700B amplifier controlled by Clampex 10.2 through Digidata 1440A data acquisition system (Molecular Devices, Sunnydale, CA, USA). Also, we stimulated different types of synapses on CA3-PCs to evaluate the influence of each synaptic input on the intrinsic excitability of CA3-PCs. Afferent PP fibers were stimulated with a concentric bipolar electrode (CBAPB125; FHC Inc., Bowdoin, ME, USA) positioned at SLM on the border of the subiculum and CA1 (for PP stimulation)(Perez-Rosello et al., 2011). Associational/commissural (A/C) fibers were stimulated with a glass micro-pipette placed on alveus. Brief stimulation pulses (0.1 ms) were generated by a computer-controlled digital stimulator (DS8000, WPI; Sarasota, FL, USA) and delivered to a stimulation electrode through an isolation unit (DLS100 stimulus isolator, WPI). For extracellular recording, field excitatory postsynaptic potentials (fEPSPs) were recorded from striatum lacunosum-moleculare (SLM) of CA3 by a glass electrode filled with 150 mM NaCl (resistance, c.a. 1 MΩ).

Prior to electrophysiological studies, an incision was made along the hippocampal sulcus of the slice to prevent possible contamination of indirect cortical input as in our previous study (Hyun et al., 2015). The stimulation intensity was adjusted so that the value of PP-EPSC was 10-20 pA in the whole cell configuration (Figs. 1, 3, 4 and 7). For extracellular recordings, the PP stimulation intensity was adjusted such that synaptic responses were lower than 20% of the maximum intensity that does not elicit population spikes (10 – 30 V, Fig. 2). For the alveus stimulation, the optimal stimulation position was found by moving the glass electrode on the slice, and the stimulation intensity was 3-5V (Fig. 2-3). For the MF stimulation (Figs. 2 and 4), we adopted minimum stimulation protocol, by which the failure rate was about 40-60% (c.a. 30 V at dentate gyrus granule cell layer; Hyun et al., 2015, Jonas 1993).

**Fig. 2.**
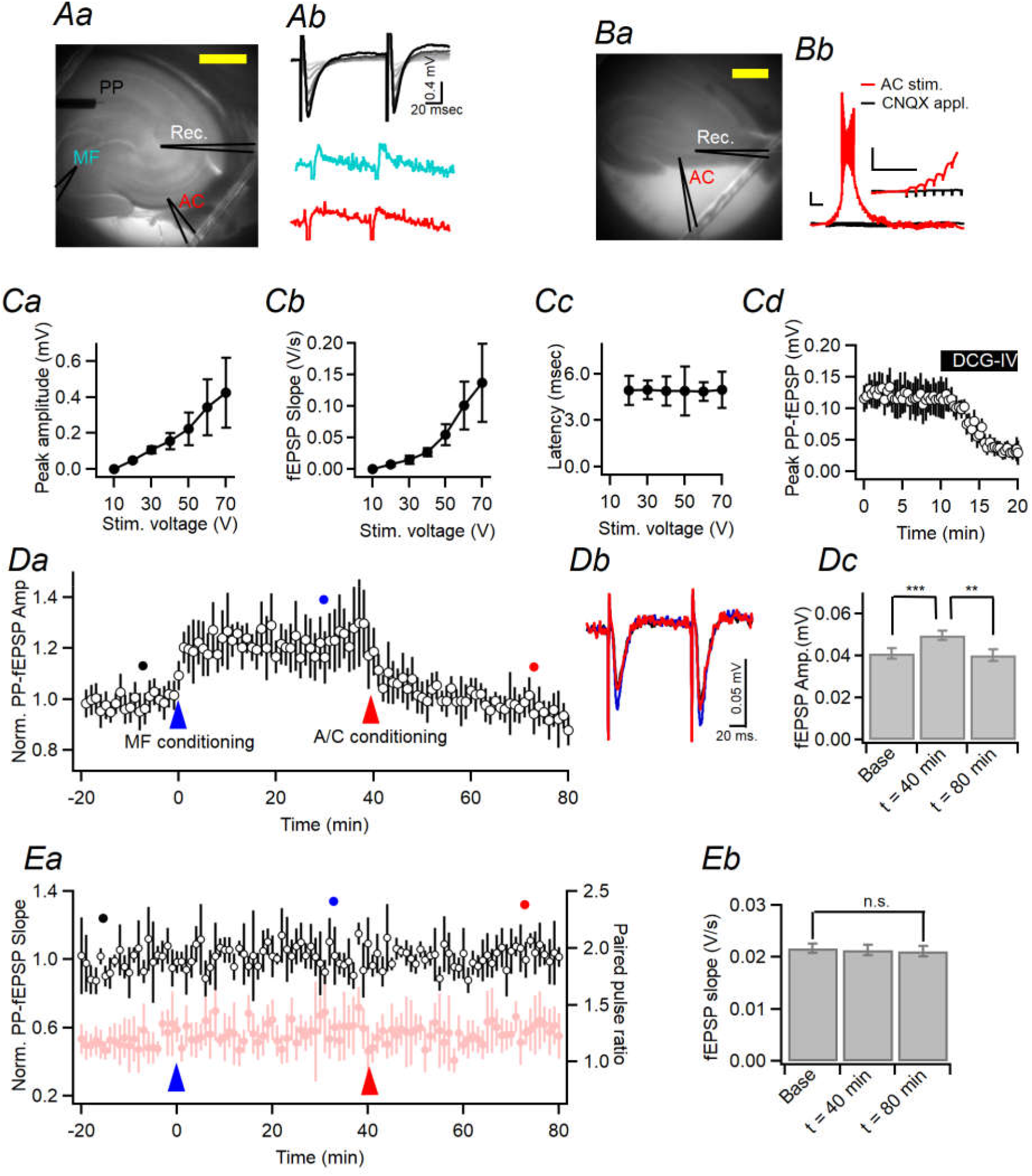
Restoration of MF-induced potentiation of PP-fEPSPs by A/C stimulation. **Aa.** IR-DIC images for extracellular recording setup. The recording electrode was placed on the CA3 SLM. Three stimulating electrodes were placed on the SLM of CA1 and subiculum border, granule cell layer of dentate gyrus, and alveus of CA3 to stimulated PP, MF, and A/C fibers, respectively. The silhouette of the each glass electrode is indicated by a black solid line. **Ab.** Upper, Representative traces of fEPSPs evoked by paired pulse stimulation of PP (PP-fEPSPs, 100 ms interval) with gradually increasing intensity ranging 10 to 70 V. Middle, MF-evoked fEPSP. Lower, A/C-evoked fEPSP trace. The same color codes were used as in Aa. **Ba.** IR-DIC images for whole-cell patch pipette on the CA3-PC and the glass electrode on the alveus of CA3 for stimulation of A/C synapses. Yellow bars in *Aa* and *Bc*, 500 μm. **Bb.** Voltage response of the CA3-PC to A/C stimulation recorded using whole-cell patch techniques before (red) and after (black) bath application of CNQX. Inset, the same EPSPs evoked by first a few stimuli in an expanded scale. Scale bars, 100 ms and 10 mV. **Ca-Cb.** Mean peak amplitude (*Ca*) and initial slope of PP-fEPSP (*Cb*) as a function of stimulation intensity. **Cc.** PP-fEPSP latency (time from PP stimulation to the peak of PP-fEPSP) was not affected by stimulus intensity (linear regression slope, t = −0.02, p = 0.98, n = 5). **Cd.** Bath application of DCG-IV attenuated the PP-fEPSP amplitudes (n = 5, t = 5.075, p = 0.007; paired t-test).**Da.** Peak amplitudes of 1st PP-fEPSP before and after MF conditioning (blue arrowhead) and A/C conditioning (red arrowhead at 40 min). The PP-fEPSPs increased and reached a plateau 5 min after MF conditioning. The PP-fEPSP stayed increased up to 40 min, and was restored to the baseline by A/C conditioning. **Db.** PP-fEPSP traces before (black; −15 min), and after MF conditioning (blue, +40 min) and after A/C conditioning (red, +80 min). Lower, the first fEPSPs in expanded time scale (gray box). The same color codes as the dots in Ca. D**c.** Summary for the peak amplitudes of PP-fEPSPs before and after MF (40 min) and A/C conditioning (80 min) (F_(2,10)_ = 17.66, p = 0.005; baseline vs. 40 min, p = 0.002; baseline vs. 80 min, p = 0.672; 40 vs. 80 min, p = 0.007; n = 6, RM-ANOVA and simple effect analysis). **Ea.** Neither the initial slope of first PP-fEPSP (black; F_(2,10)_ = 0.174, p = 0.842) nor paired pulse ratio (PPR, light red; F_(2,10)_ = 0.057, p = 0.944, RM-ANOVA and simple effect analysis) measured from the peak amplitudes of 1^st^ and 2^nd^ PP-fEPSPs were affected by the MF and A/C conditioning. **Eb.** Summary for the initial slope of PP-fEPSPs at the baseline, after MF conditioning and after A/C conditioning.\

**Fig. 3.**
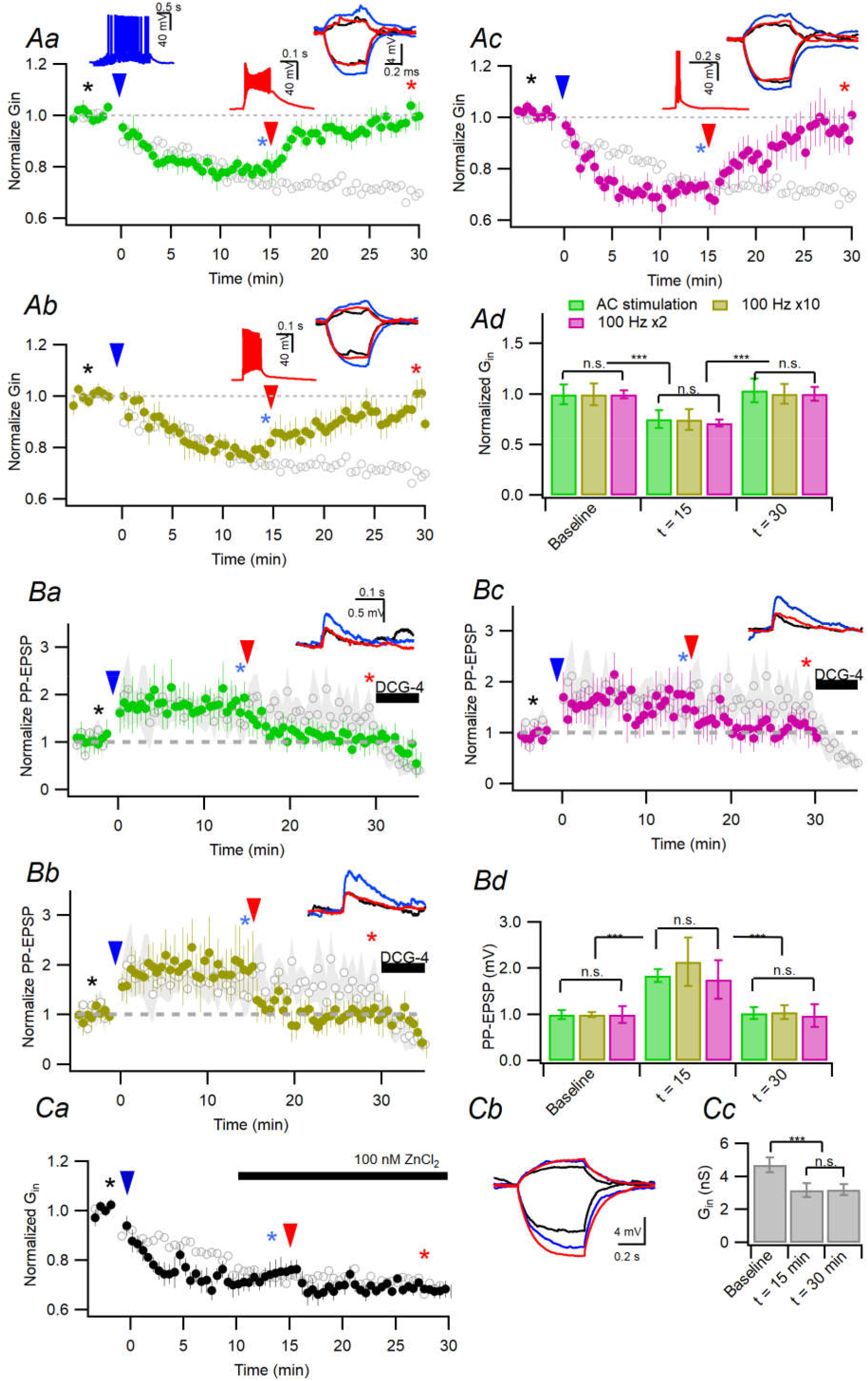
Depotentiation of MF-induced LTP-IE can be induced by somatic or A/C synaptic stimulation, and is hindered by Zn^2+^. **Aa-Ac**. Baseline- normalized G_in_ (filled circles) changes caused by MF conditioning at 0 min (blue arrowhead) and by somatic AP bursts at 15 min (red arrowhead) evoked by A/C stimulation (*Aa*) or direct somatic current injection of ten (*Ab*) or two (*Ac*) pulses at 100 Hz. For A/C stimulation (*Aa*), 0 vs. 15 min, p < 0.001; 0 vs. 30 min, p = 0.945; 15 vs. 30 min, p < 0.001 (n = 7, GLM and simple effect analysis). For somatic AP bursts (*Ab*), 0 vs. 15 min, p < 0.001; 0 vs. 30 min, p = 0.936; 15 vs. 30 min, p < 0.001 (n = 7, GLM and simple effect analysis). For two AP simulation (*Ac*), 0 vs. 15 min, p < 0.001; 0 vs. 30 min, p = 0.439; 15 vs. 30 min, p < 0.001 (n = 7, GLM and simple effect analysis). The MF-induced decrease of G_in_ in the absence of intracellular GSH is reproduced from (Eom et al., 2019) for comparison (*gray symbols*). For A/C synaptic activation, CA3 alveus was stimulated at 20 Hz for 0.5 s (A/C conditioning, Aa). *Left inset*, Voltage response to MF conditioning. Middle inset, AP bursts induced by A/C conditioning (Aa) or somatic current injection (Ab and Ac). *Right inset*, Representative voltage responses to somatic subthreshold current injection (+10 and −30 pA) for measuring G_in_ at the time indicated by black (baseline), blue (at 14 min), and red (at 29 min) asterisks. The same color codes were used for the asterisks and traces. **Ad.** Summary for normalized G_in_ at the baseline, 15 min after MF conditioning, 15 min after the depotentiation (time: F_(2,18)_ = 162.345, p < 0.001, stimuli: F_(2,18)_ = 1.392, p = 0.274, stimuli × time: F_(2,18)_ = 1.66, p = 0.218; GLM and simple effect analysis). **Ba-c.** Same as Aa-Ac but peak amplitudes of PP-EPSP were monitored instead of G_in_. For alveus stimulation (*Ba*), 0 vs. 15 min: p = 0.012; 0 vs. 30 min, p = 0.818, 15 vs. 30 min, p = 0.011 (n = 7, GLM and simple effect analysis). For 10 somatic APs (*Bb*), 0 vs. 15 min, p = 0.012; 0 vs. 30 min, p = 0.848; 15 vs. 30 min, p = 0.011 (n = 7, GLM and simple effect analysis). For 2 Aps (*Bc*), 0 vs. 15 min, p = 0.006; 0 vs. 30 min, p = 0.801; 15 vs. 30 min, p = 0.004 (n = 7, GLM and simple effect analysis). *Gray symbols*, PP-EPSP amplitudes measured without intracellular GSH, which is reproduced from (Eom et al., 2019) for comparison. *Blue arrowheads*, MF conditioning at 0 min. *Read arrowheads*, second conditioning by CA3 alveus stimulation (*Ba*) or somatic injection of current pulses (*Bb* and *Bc*) at 15 min. *Insets*, Representative traces for PP-EPSPs recorded at time points indicated by asterisks with the same color codes. **Bd**. Summary for the PP-EPSP amplitudes at baseline, 15 min after MF conditioning, 15 min after the second conditioning (time: F_(2,18)_ = 71.199, p < 0.001, stimuli: F_(2,18)_ = 1.392, p = 0.274, stimuli ;× time: F_(2,18)_ = 0.822, p = 0.520; GLM and simple effect analysis). **Ca.** Time course of normalized G_in_ changes (filled circles) caused by MF conditioning (20 Hz for 2 s) at 0 min (blue arrowhead) and subsequent somatic 100 Hz AP bursts for 0.1 s at 15 min (red arrowhead). For comparison, MF-induced decrease of G_in_ measured in the absence of intracellular GSH is reproduced from previous study (Eom et al., 2019) (gray symbols). ZnCl_2_ was bath-applied 10 min after MF conditioning. Note that somatic AP bursts did not trigger depotentiation of MF-induced LTP-IE in the presence of 100 nM ZnCl_2_. (15 min vs. 30 min, p = 1.00; simple effect analysis). **Cb.** Representative voltage traces in response to subthreshold current (+10 and −30 pA) for measuring G_in_. Voltage responses were measured before and c.a. 15 min after MF conditioning, and c.a. 15 min after somatic AP bursts (indicated by black, blue and red asterisks in Aa, respectively). **Cc.** Summary for the mean G_in_ at baseline, 15 min after MF conditioning, and 15 min after 100 Hz AP bursts (F_(1,4)_ = 16.109, p = 0.016, RM-ANOVA; 0 vs. 15 min, p = 0.007; 0 vs. 30 min, p = 0.002; 15 vs. 30 min, p = 1.00; n = 5, GLM and simple effect analysis).

**Fig. 4.**
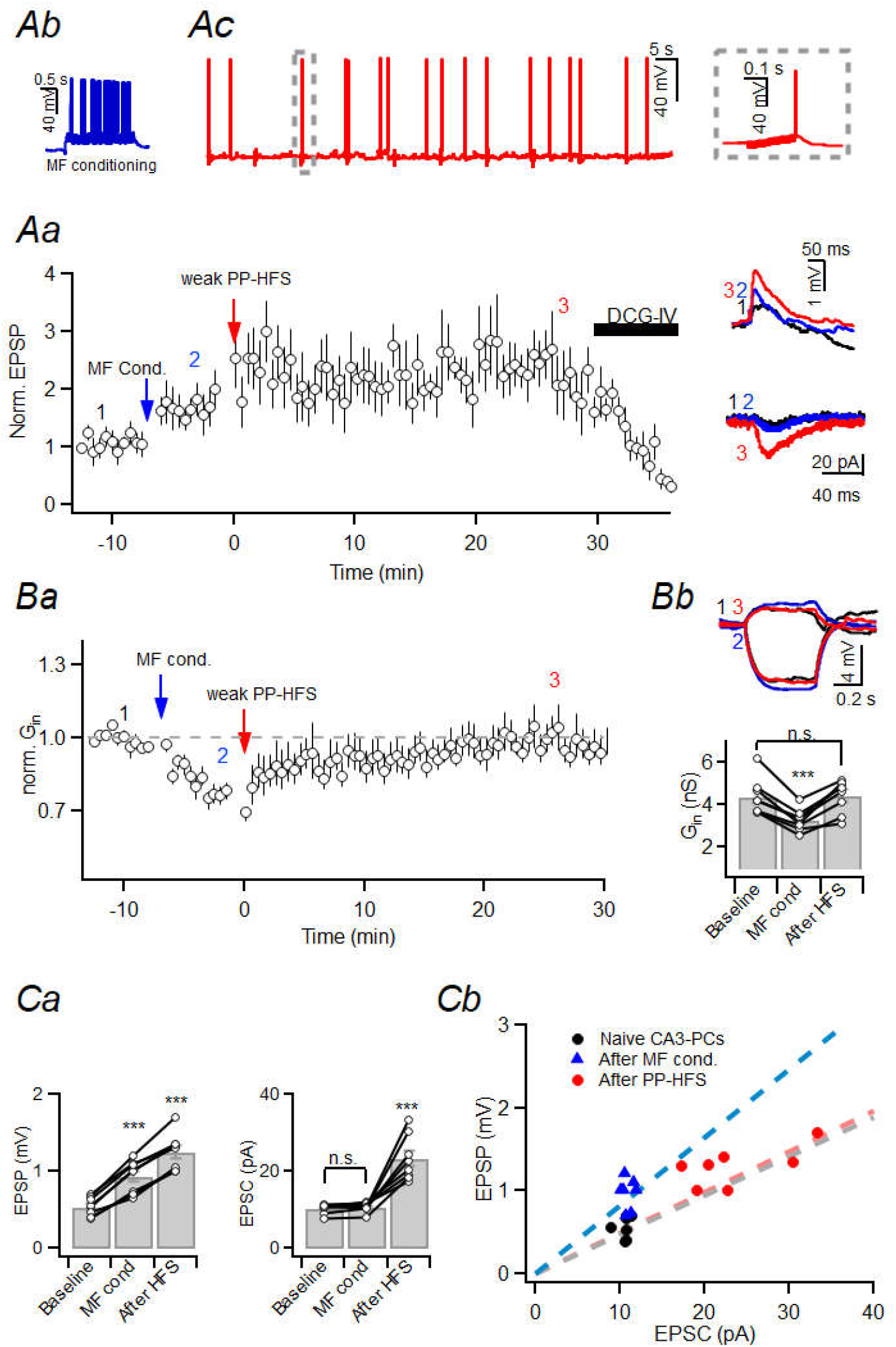
MF-induced LTP-IE is terminated after homosynaptic PP-LTP. WCRs of CA3-PCs was done using GSH-containing pipette solution. **Aa.** Baseline-normalized amplitudes of PP-EPSPs were increased by MF conditioning (20 Hz MF stimulation for 2 s, *blue arrow*) and further by high-frequency stimulation of PP (PP-HFS) with weak stimulation intensity (*red arrow*). *Insets*, Representative traces of PP-EPSPs (upper) and PP-EPSCs (lower) and before (*black*), after MF conditioning (*blue*), and after PP-HFS (*red*). The recording time points are indicated by numbers in the main graph. **Ab.** Representative traces for voltage response to MF conditioning. **Ac.** Voltage response to weak PP-HFS (10 bursts every 10 s, 20 pulses at 100 Hz per a burst; baseline EPSC < 12.55 pA). Inset, gray boxed region in an expanded time scale. **Ba.** Baseline-normalized G_in_ was decreased by MF conditioning (*blue arrow*) and restored by PP-HFS (*red arrow*). G_in_ was monitored in the same cells as in Aa (F_(1,4)_ = 24.064, p < 0.001; RM-ANOVA). **Bb**. *Upper*, Representative voltage responses to somatic current injection of +10 and −30 pA at baseline (black), after MF conditioning (blue), and after PP-HFS (red). *Lower*, Mean G_in_ at baseline, 6 min after MF conditioning, and 25 min after PP-HFS. Baseline vs. post-MF conditioning, p < 0.001; baseline vs post-PPHFS, p = 1.00; post-cond. vs post-PPHFS, p < 0.001 (n = 7, GLM and simple effect analysis). **Ca.** Summary for the mean amplitudes of PP-EPSPs (left, F_(1,4)_ = 138.09, p < 0.001; RM-ANOVA) and PP-EPSCs (right, F_(1,4)_ = 35.751, p < 0.001; RM-ANOVA) at baseline, after MF conditioning, and after PP-HFS. For EPSP, baseline vs. post-MF cond., p < 0.001; baseline vs. post-PPHFS, p < 0.001; post-MF cond. vs. post-PPHFS, p < 0.001; (n = 7, GLM and simple effect analysis). For EPSCs, baseline vs. post-MF cond., p = 0.647; baseline vs. post-PPHFS, p < 0.001; post-MF cond. vs post-PPHFS, p < 0.001 (n = 7, GLM and simple effect analysis); **Cb.** Scatterplot for PP-EPSP vs. PP-EPSC measured in the same CA3-PCs at baseline (black), after MF conditioning (blue), and after PP-HFS (red). Baseline vs. post-cond., p < 0.001; baseline vs post-HFS, p = 1.00; post-cond. vs post-HFS, p < 0.001; (n = 7, GLM and simple effect analysis).

### Stereotaxic surgery and intrahippocampal injection of adeno-associated virus (AAV) for ex vivo study

An adeno-associated virus (AAV-RAM-d2TTATRE-EGFP-WPRE-pA, titer: 4.8 × 10^12^ GC/ml) was manufactured by KIST virus facility (Seoul, Republic of Korea). Surgery was performed on young-adult SD rats (6 weeks). Before stereotaxic injection of the AAV into hippocampus, animals were singly housed to new homecage for 1 week to acclimation with doxycycline diet (40 mg/kg). Animals were kept on the doxycycline diet and allowed to recover for at least 7 days following surgery. After anesthesia with isoflurane, rats were secured in a stereotaxic frame (Neurostar, Tubingen, Germany). Holes were drilled bilaterally in the skull at the injection sites (2 sites). Stereotaxic coordinates used for intrahippocampal injections were as follows (from bregma); for a ventral hippocampus, anterior–posterior −2.85 mm, mediolateral ±2.8 mm, dorsoventral 3.3 mm. A 33-gauge needle attached to a 50 μl Hamilton syringe (#80908), mounted to the stereotaxic frame, and under control of a microinjection syringe pump UMP3T-1 controller (WPI, Sarasota, FL, USA) was used to inject 0.8 μl of AAV at each site. Injections occurred at a rate of 0.08 μl/min, after which the needle was left in place for an additional 2 min. After injections were completed, the skin was sutured and the animals were allowed to recover for 1 h on a heating pad before returning to the home cage. Mice remained in the homecage for an additional 2 weeks to recover before the start of behavioral testing and post-hoc *ex vivo* experiments. The diet was switched to doxycycline-free diet 2 days before contextual fear conditioning (CFC) to allow tetracycline transcriptional activator (tTA) to activate the tTA-responsive element (TRE) promoter driving expression of EGFP.

### B6.Cg-Tg(Fos-tTA,Fos-EGFP*)1Mmay/J mice

cfos-tTA/cfos-shEGFP mice were purchased from Jackson laboratory (stock No: 018306; Donating investigator: Mark Mayford, The Scripps Research Institute, La Jolla, CA), and crossed to the C57BL/6J background to maintain lineage. The cfos-shEGFP transgenes allows us to quantify c-fos expression correlated with neuronal activation by measuring fluorescence of short half-life green fluorescent protein (shEGFP, t_1/2_ = 2 hours). For genotyping, DNA was isolated from the tail of each mouse in the litter aged 6 ∼ 8 days as the method provided by the Jackson Laboratory (https://www.jax.org/strain/018306). All animals were maintained in the standard environmental conditions described above.

### Exposure of animals to a novel context

The cfos-tTA/cfos-shEGFP mice (hereafter cfos-EGFP mice) of either sex (6-8 weeks old) were singly housed for 1 weeks before the experiments. On the experimental day, the mice were placed in novel contexts; context A or A and B. Context A is a chamber (18 cm wide × 18 cm long × 30 cm high; H10-11M-TC; Coulbourn Instruments 5583, PA 18052, USA) consisting of a metal grid floor, aluminum side walls, and a clear Plexiglass front door and back wall. The context A chamber was lit indirectly with a 12 W light bulb. The features of context B were the same as context A, except for a unique odor (1% acetic acid), dimmer light (50% of A), and a slanted floor by 15° angle. The chamber and cage were cleaned using 70% ethanol before placing an animal and between each runs. In context A, but not in context B, animals were allowed a 3-min exploration, received a single foot shock (0.7 mA, for 2 s) and were returned to their home cage 60 s after the shock. In the context B, animals were allowed a 5-min exploration and returned to their homecage.

### Confocal microscopy and image quantification

We tested 4 cfos-shEGFP mice (PW6-8) between 8 and 14 weeks of age for contextual exposure. Before the experiments, mice were a singly housed to new homecage for 1 week for acclimation. For experiments depicted in Fig. S4, mice were divided into two groups: exposure of context A only and waiting in homecage for 90 min (ctxA), exposure of context A and B with 25-min interval and waiting in homecage for 60 min (ctxAB). Two groups were sacrificed to prepare hippocampal slices after 90 min from the exposure of ctxA, as aforementioned. Confocal images from the hippocampal slices in Fig. S4 were obtained in the submerged recording chamber that was continuously perfused with the recording solution at 1.5 ml/min. To suppress unnecessary neural activity, picrotoxin (10 μM) and CNQX (10 μM) were added to the solution. For analysis of fluorescence intensity of shEGFP(+) CA3-PCs, Z-series (60-90 frames, 0.71 μm /frame) image were acquired from the live slices using a 40x lens using FV 1200 confocal laser scanning microscope (Olympus, Tokyo, Japan). LD473 (15mW) laser system controlled by combiner (FV10-MCPSU) was used to excite shEGFP in CA3-PCs. Typically, four images were analyzed for each animal in the CA3 region. Fluorescence intensity profile for each shEGFP(+) CA3-PCs based on depth from surface of slices were obtained from Z-stack image of CA3 region. The surface of hippocampal slices was defined as 0 μm of depth in hippocampal slices. We regarded fluorescence signals that is above a threshold of 5× the SD of baseline fluctuation of signal intensity profile as the signal of shEGFP(+) CA3-PCs. The depth for the maximal fluorescence intensity was defined as the depth of CA3-PCs.

### Epifluorescence imaging for AAV-injected rats or cfos-shEGFP mice

Fig. 5 and 6 ∼ 8 depict ex vivo electrophysiological studies of CA3-PCs obtained from AAV-injected rats and cfos-shEGFP mice, respectively. After acclimation of 1 weeks, AAV-injected rats were exposed to context A once for 5 min and returned to homecage for 1, 1.5, and 3 hours (Fig. 5) and sacrificed to obtain brain slices. cfos-shEGFP mice were divided into several groups: exposure of context A only and waiting in homecage for 60 min (ctxA60), exposure of context A only and waiting in homecage for 90 min (ctxA90), and exposure of context A and B with 25-min interval and waiting in homecage for 60 min (ctxAB). ctxA60 and ctxA90 groups were sacrificed to prepare brain slices after 60 min and 90 min from the exposure of context A. For ctxAB group, animals were sacrificed to obtain brain slices after 60 min from the exploration of last context, context B. EGFP or shEGFP was excited at 488 nm light with a monochromator (Polychrome IV, xenon-lamp based, Polychrome-V, TILL-Photonics, Martinsried, Germany) and visualized using a 520/535-nm band-pass emission filter (Fig. 5 and 6). In experiments for Fig. 6–8, to adjust deviation of fluorescence intensity of shEGFP caused by the optical path length, 50 μM Alexa flour 594 (AF594) was added to intracellular solution and loaded to CA3-PCs during whole-cell configuration. AF594 was excited at 594 nm light with the monochromator and visualized using a 660 nm long-pass emission filter. Acquired images were analyzed with HC-Image Live 4.0 (Hamamatsu Photonics, Shizuoka, Japan). Whole-cell recordings (WCR) were performed under the IR-DIC (Fig. 5Ab). WCR of EGFP(-) cells were used as control and compared to recording of EGFP(+) or shEGFP(+) cells. The fluorescence intensity in the wide-field epifluorescence image was set with manually-adjusted ROI based on the contour of cells.

**Fig. 5.**
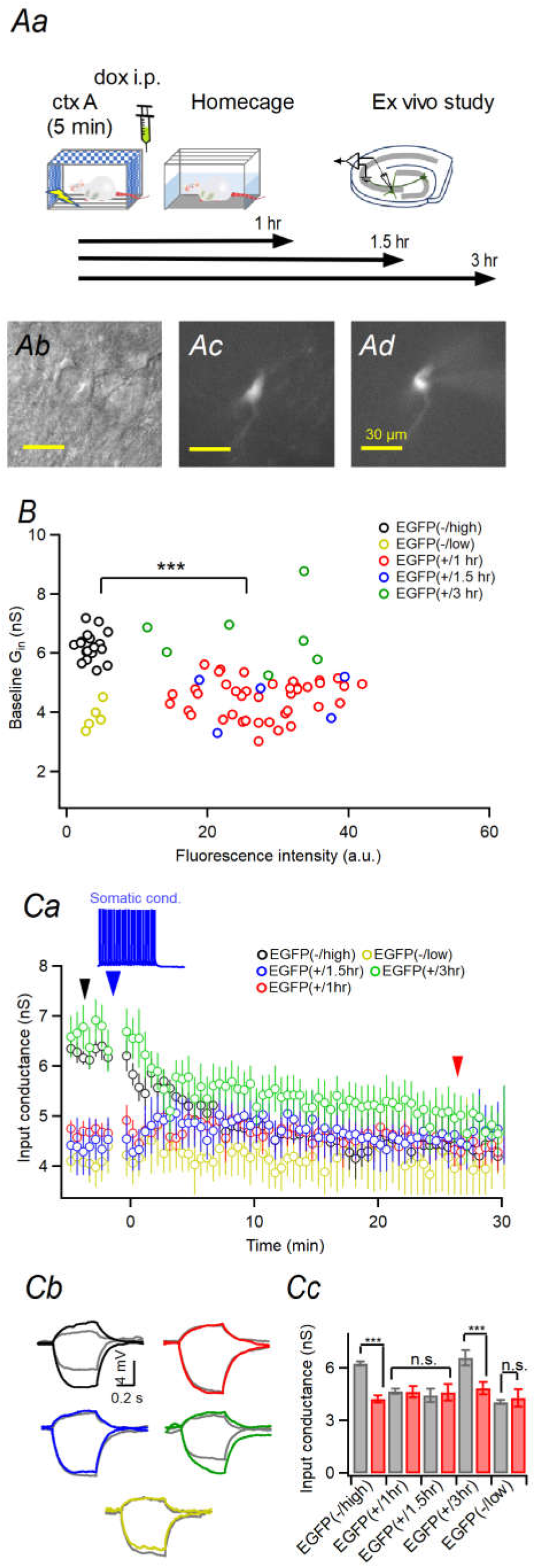
The LTP-IE of CA3-PC did not last longer than 3 hours in vivo. **Aa.** Schematic timeline of the experimental procedure. Doxycycline was administrated via intraperitoneal route to prevent further expression of GFP after visiting context A. Animals were sacrificed to prepare hippocampal slices after 1 hr, 1.5 hrs, and 3 hrs. **Ac-Ad.** IR-DIC image (*Ab*) and fluorescence image excited at 488 nm for GFP(+) CA3- PCs (*Ac*: immediately after making whole-cell configuration; *Ad*: 30 min after whole-cell configuration. Note that diffusion of GFP to the whole cell pipette in *Ad*). Scalebar, 30 μm. **B.** Baseline G_in_ as a function of GFP fluorescence intensity (F_EGFP_). Most GFP(-) cells (F_EGFP_ = 3.38 ± 0.30 a.u.) showed high baseline G_in_ [denoted as GFP(-/high) cells]. Some of GFP(-) cells (3.97 ± 0.47 a.u.), however, showed lower baseline G_in_ [denoted as GFP(-/low) cells]. GFP(+) cells after 1 hr (27.68 ± 1.11 a.u.) and 1.5 hr (28.92 ± 4.16 a.u) showed lower G_in_. In contrast, GFP(+) cells after 3 hr (25.72 ± 3.63 a.u.) showed high G_in_ similar to GFP(-/high) cells. **Ca.** Time course of G_in_ before and after somatic conditioning (indicated as blue arrowhead). Time points at which G_in_ was measured for comparison between cell groups and the representative traces are shown in *Cb* are indicated as black and red arrowheads (before and 30 min after somatic conditioning, respectively). **Cb-Cc.** Representative voltage traces for different cell groups (same color code as *B* and *Ca*). Traces for baseline G_in_ [GFP(-/high), 6.25 ± 0.26 nS (n = 8); GFP(+) at 1 hr, 4.68 ± 0.33 nS (n = 8); GFP(+) at 1.5 hr, 4.48 ± 0.33 nS (n = 5); GFP(+) at 3 hr, 6.59 ± 0.28 nS, p < 0.001 (n = 7)] were indicated as gray color. **Cc**, Summary of G_in_ before (gray) and after (red) somatic conditioning (time, F_(1,24)_ = 66.042; fluorescence, F_(3,24)_ = 3.705, p = 0.025; time x fluorescence, F_(3,24)_ = 25.712, p < 0.001, RM-ANOVA; For 1 hr, baseline vs. post-SC, p = 0.837; For 1.5 hr, baseline vs. post-SC, p =0.554; GLM and simple effect analysis).

**Fig. 6.**
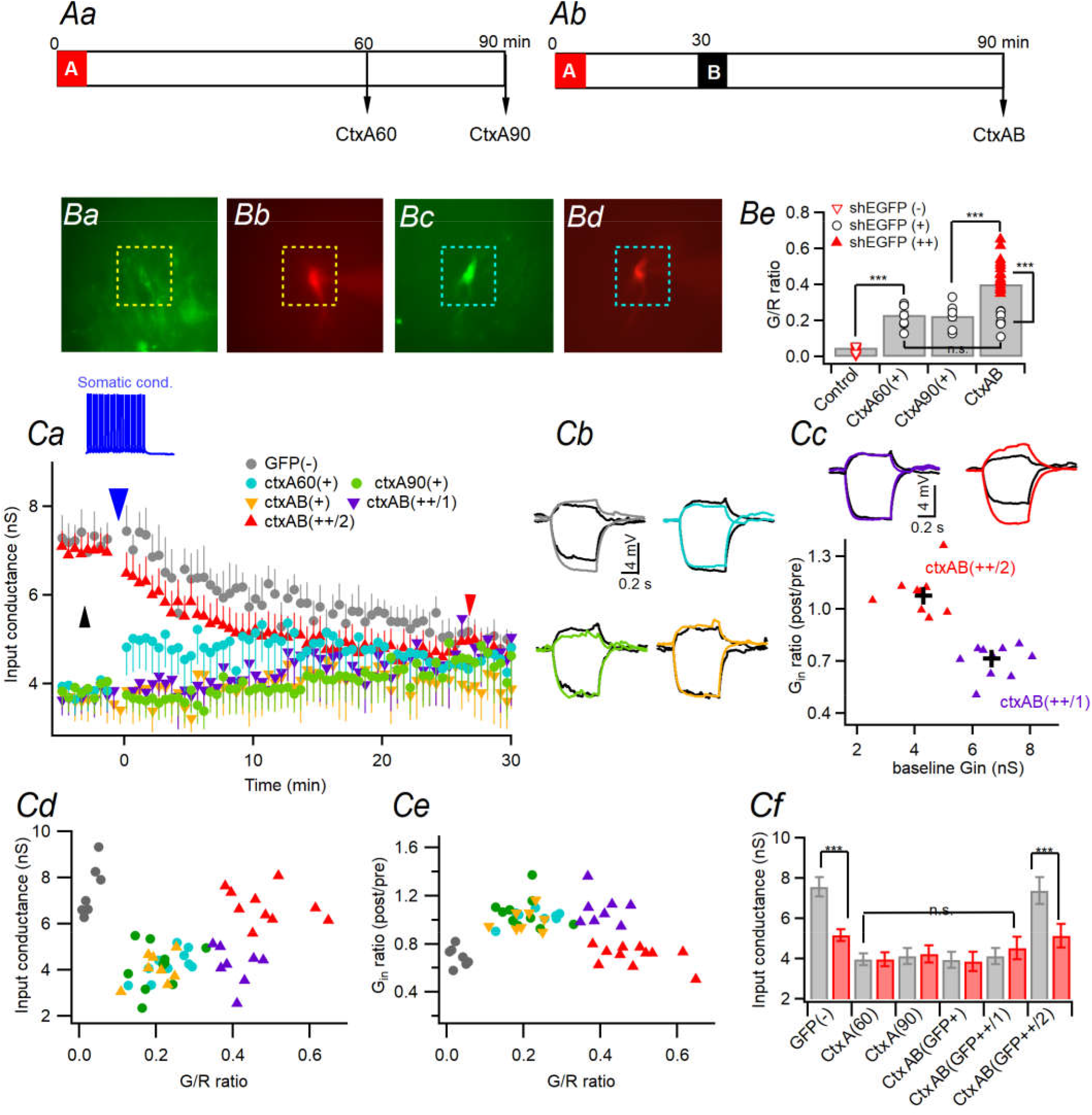
LTP-IE occurs in CA3 ensemble cells and may be reversed in part of twice-activated ensemble cells. **Aa.** Timeline of experimental procedures. *Upper*: Mice were subject to CFC in context A for 5 min, and kept in homecage for 1 hr or 1.5 hr before ex vivo study (ctxA60 and ctxA90 mice). **Ab**. Mice additionally visited context B 25 min after CFC, and then were kept in homecage (ctxAB mice). **Ba-d.** Fluorescence images of two AF594-loaded CA3-PCs with low (Ba-b) and high (Bc-d) G/R ratios. **Be.** Summary for G/R ratios. **Ca.** Time course of G_in_ changes in response to somatic conditioning. Somatic conditioning (SC) induced LTP-IE in GFP(-) CA3-PCs (gray symbols; n = 6, pre- vs. post-SC G_in_, p < 0.001), but not other cells except ctxAB(++/2) cells. *Inset*, voltage response to 10 Hz somatic conditioning for 2 s (blue arrowhead). Small arrowheads indicate the time at which representative traces shown in *Cb* and *Cc* were recorded. **Cb.** Representative traces for voltage responses to −30 pA and +10 pA current steps in GFP(-), ctxA60(+), ctxA90(+), and ctxAB(+) cells. Traces before conditioning are depicted in gray color, and those after somatic conditioning are in the same color codes as in *Ca*. **Cc.** *Upper*, Representative voltage traces for G_in_ in ctxAB(++/1) and ctxAB(++/2) cells. *Lower*: ctxAB(++) cells were clustered into two groups based on baseline G_in_ and G_in_ ratios before and after somatic conditioning using k-means analysis. There was a significant difference between CtxAB(++/1) and CtxAB(++/2) groups, and the centroids of two cell groups were depicted as cross symbols. **Cd-Ce.** Baseline G_in_ (*Cd*) and G_in_ ratio (*Ce*) as a function of green-to-red fluorescence ratio (G/R ratio). The same color codes used in *Ca*-*Ce*.

**Fig. 7.**
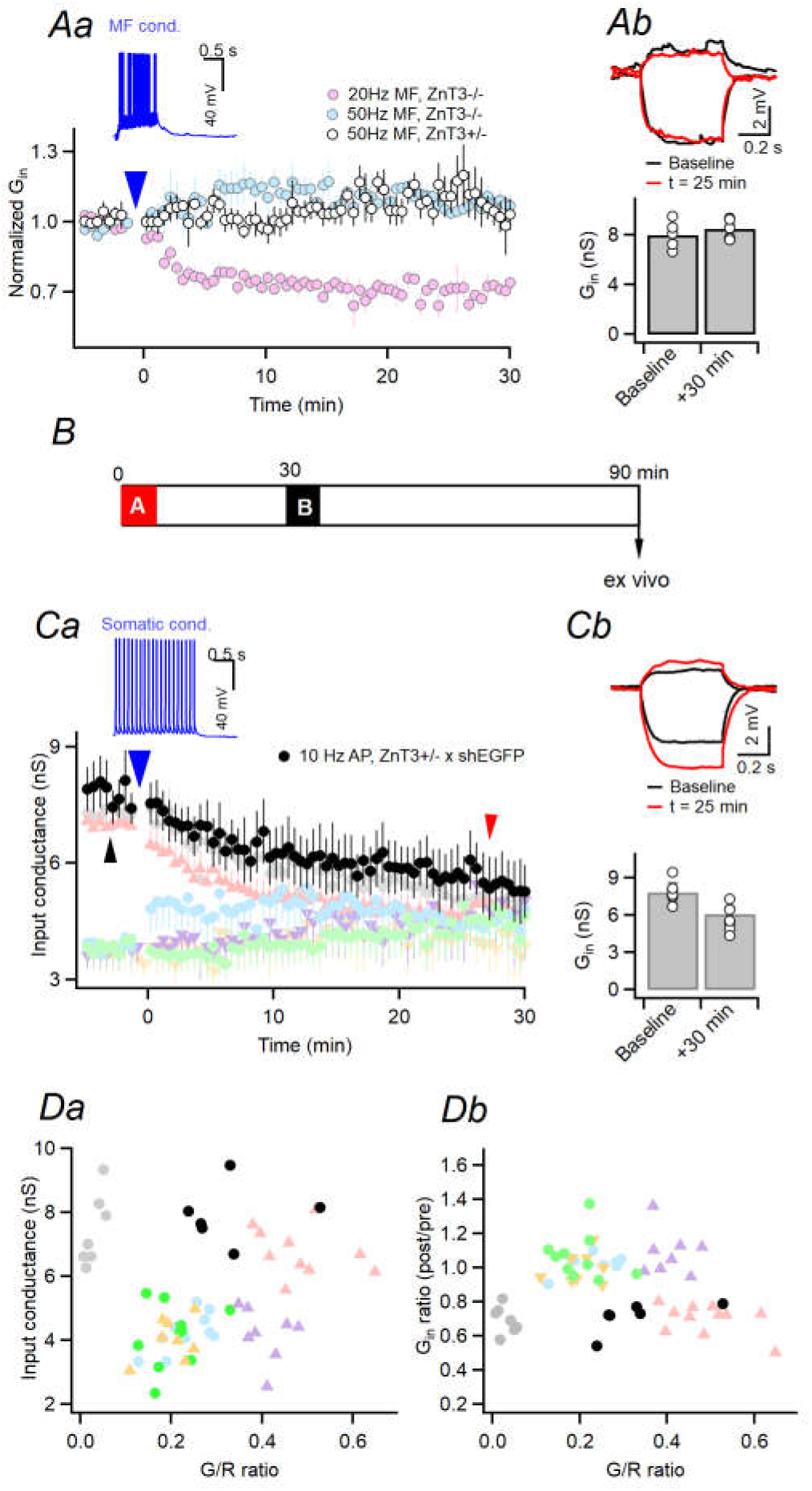
LTP-IE does not occur in CA3 ensemble cells of ZnT3 hetero-knockout mice. **Aa.** G_in_ changes before and after 50 Hz MF stimulation for 1 s in CA3-PCs of ZnT3HT mice (open symbols). The G_in_ changes induced by 20 Hz 2 s MF (pink) and 50 Hz 1s MF stimulation (light blue) were reproduced from our previous study (Eom et al. 2019). **Ab.** *Upper*: Representative traces for G_in_ before (black) and 25 min after MF conditioning (upper) in CA3-PCs of ZnT3HT mice. *Lower*: Summary for G_in_ before and after 50 Hz MF conditioning in CA3-PCs from ZnT3HT mice (n = 5, t = −1.141, p = 0.318, paired t-test). **B.** Timeline of experimental procedure. **Ca.** G_in_ before and after 10 Hz 2 s somatic conditioning in CA3-PCs of ZnT3-HT/cfos-shEGFP mice (black),. G_in_ before and after somatic conditioning in other groups of CA3-PCs were reproduced from previous figure (Fig. 6, the same but pale colour codes). **Cb.** Representative traces for G_in_ before (black) and 25 min after conditioning (*upper*) and summary of G_in_ before and after somatic conditioning in CA3- PCs from ZnT3HT/cfos-shEGFP mice (n = 6, t = 7.675, p = 0.001, paired t-test). **Da-b.** Baseline G_in_ (*Ca*) and post/pre-conditioning G_in_ ratio (*Cb*) as a function of G/R ratio in CA3-PCs of ZnT3-HT/cfos-shEGFP mice (black). Colored symbols are reproduced from Fig. 6 with pale color.

**Fig. 8.**
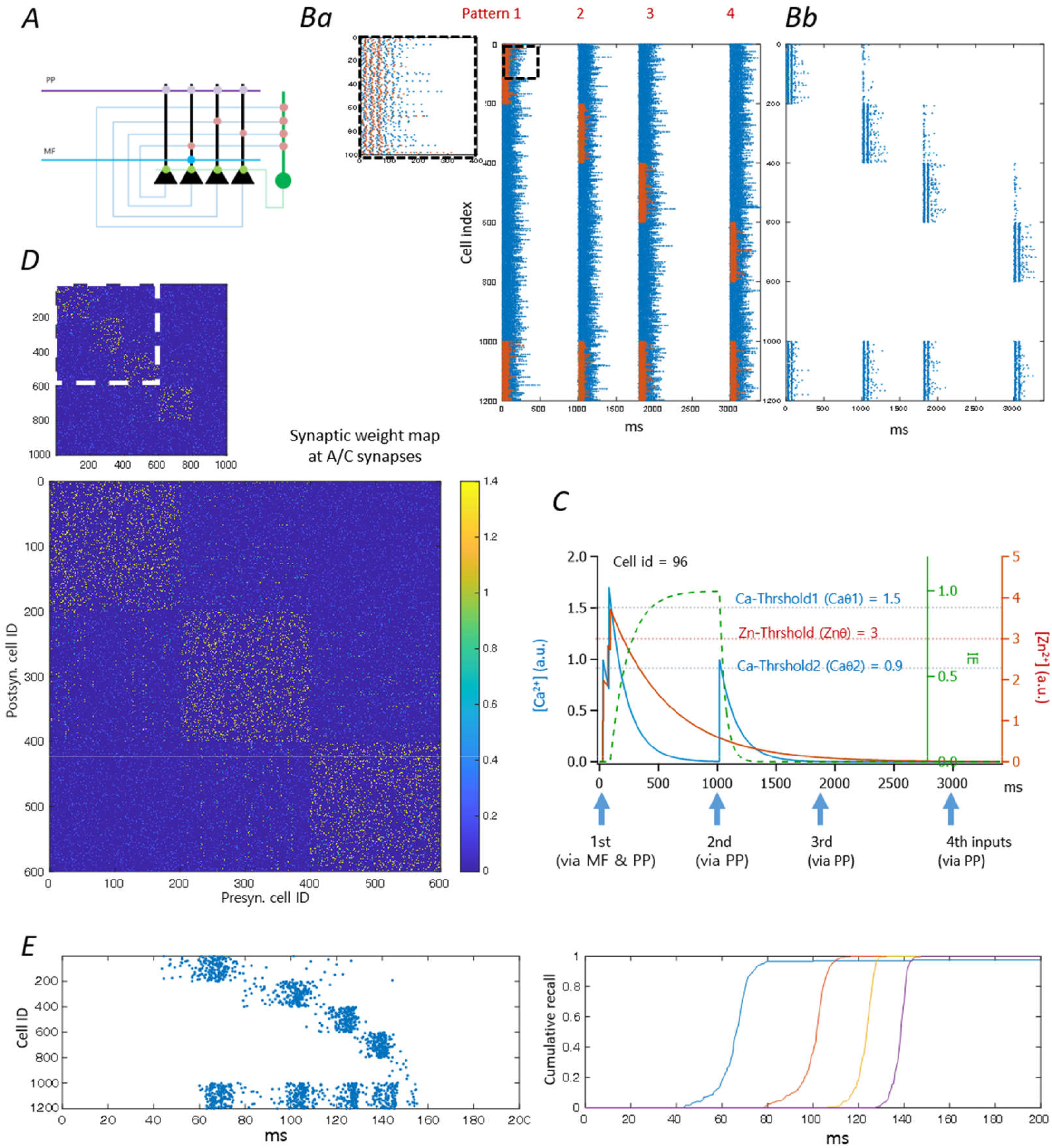
Input-specific bi-directional regulation of intrinsic excitability may contribute to sequential memories. **A.** Schematic illustration of CA3 network model comprised of pyramidal cells (PCs, black triangles) and an inhibitory interneuron (IN, green circle). **Ba.** Input patterns via MF (red dots) and PP (blue dots) during encoding phase. Inset, broken boxed region in an expanded scale. **Bb**. Raster plot of network responses to the input patterns shown in Ba. The distributions for the number of MF and PP inputs per a pattern are shown in Fig. S7D. The cell indices of 1 to 1000 are PCs, and the rest cell indices (1001 ∼ 1200) are INs. **C,** Ca^2+^ (blue) and Zn^2+^ (red) transients in a PC in the **M_1_** subset (cell id = 96) during the encoding phase. The timings of input pattern arrival are indicated by arrows below the abscissa. The intrinsic excitability index (IE, green broken line) starts to increase when both [Ca^2+^] and [Zn^2+^] are higher than their thresholds (Caθ1 = 1.5 a.u.; Znθ = 3 a.u.). When IE > 0.7, the arrival of subsequent PP inputs elicits a CaT devoid of Zn^2+^ signaling. When CaTs is higher than Caθ2 (= 0.9 a.u.) without Zn^2+^ signal, the IE returns to the baseline. Afterwards, this PC does not fire any longer in response to PP inputs alone. Note that the kinetics of Ca^2+^, Zn^2+^ and IE were assumed faster than real values for the sake of reducing computing time (decay τ for CaT and ZnT = 150 and 300 ms, respectively; rise and decay τ for IE = 150 and 50 ms, respectively). **D.** Part of matrix of A/C synaptic weights (boxed in the Inset) resulting from training the recurrent network using a series of 4 patterns shown in Ba. Rows and columns represent indices of post- and pre-synaptic CA3-PCs, respectively. Synaptic weights are color-coded as shown on the right bar. Note that sparse potentiation of synaptic weights between **M_1_** (cell index, 1-200) and **M_2_** (cell index, 201-400) but not between **M_1_** and **M_3_** (cell index, 401-600). **E**, *Left*, Raster plot of network responses to the PP inputs shown in Fig. S7E. Note that most PP inputs are terminated in 30 ms (Fig. S7E), but the complete recall of **M_1_** is accomplished by recurrent connections that have been strengthened during encoding phase. In turn, hetero-association between **M_1_** and **M_2_** that occurred during encoding phase allows subsequent activation of **M_2_**, and so on. Right, cumulative fraction of activated PCs in each ensemble.

### Simulation methods

We considered a recurrent network comprised of 1000 PCs and 200 INs, in which PCs and INs are numbered with indices from 1 to 1000, and those from 1001 to 1200, respectively. Initial connectivity matrix (*W*) are shown in Fig. S7A. The connectivity, defined as the probability that an element of *W* has nonzero value, was set to 10% for synaptic connections between PCs, while the connectivity between CA3 PCs is less than 2% *in vivo* (Guzman et al., 2016). This high connectivity partially compensates lower recurrent synaptic inputs from a given ensemble compared to the real rat CA3 network. The firing property of neurons was simulated by the leaky integrate and fire model as described in detail elsewhere (Yoon et al., 2020). The resting membrane potential (V_L_) and the firing threshold (V_θ_) were set to −65 and −55 mV, respectively, for both CA3-PCs and GABAergic interneurons (INs). To model recurrent and inhibitory synaptic inputs, we assumed that firing of a presynaptic CA3 PC or IN (denoted by subscripts ‘e’ and ‘i’, respectively) induces an instantaneous increase in the postsynaptic conductance by G_s_ after a synaptic delay, D_s_, where s ∈ {e, i} (D_e_ = 2 ms; D_i_ = 1 ms), and that the decay of the conductance change follows a mono-exponential function with a time constant τ_s_ (τ_e_ = τ_i_ = 10 ms) (Guzman et al., 2016; Kohus et al., 2016). Multiple synaptic inputs to a neuron, *j*, were linearly summated. When the number of *s* type presynaptic neurons innervating postsynaptic neuron, *j*, is K_s_, the dynamics of subthreshold membrane potential of the neuron, *j*, is assumed to follow the equations:

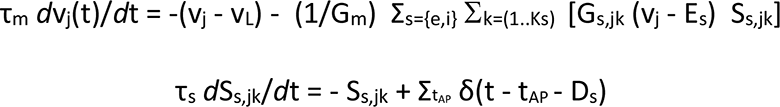

where τ_m_ = cell membrane time constant; G_m_ = passive membrane conductance (5 nS for PCs and INs); G_s, jk_ = synaptic conductance of synaptic type, *s*, from a neuron *k* to *j*; E_s_ = reversal potential of synapse type (E_e_ = 0 mV, E_i_ = −68 mV). The *S*_s,jk_ is an activation parameter for synaptic type *s* from a neuron *k* to *j*. It is activated by presynaptic firing at t_AP_, and undergoes mono-exponential decays with a time constant, τ_s_. When membrane potential (*v*) surpasses the threshold potential (V_θ_), the *v* was reset to −55 mV. To simulate a post-spike refractory period, the V_θ_ was increased to −15 mV just after firing, which undergoes mono-exponential decay with time constants of 50 and 15 ms for PC and INs, respectively. The summation of external excitatory synaptic inputs via MF and PP were calculated in the same manner as recurrent excitatory inputs, but the arrival of external synaptic inputs was pre-determined as a patterned input as shown in Fig. 8Ba. We assumed that four input patterns arrive at t = 0, 0.8 s, 1.8 s and 3 s in a series. The synaptic conductance values for MF and PP inputs to PCs were set to 20 and 0.3 nS, and those to INs were 4 and 0.15 nS, respectively. The decay time constants of MF and PP inputs were set to 4 and 17 ms (Hyun et al., 2015; Jonas et al., 1993). At the start of simulation, membrane potential of all neurons was initialized to −65 mV. The voltage derivative (dv/dt) of each neuron was calculated with a time step of 0.4 ms, and was numerically integrated by Euler methods using Matlab software (R2021a, Mathworks, USA). During encoding phase, the recurrent and PP synaptic weights were modified under the STDP rules. The dependence of LTP level on the spike timing difference between pre and post-synaptic cells (Δt = t_post_ – t_pre_) is shown in Fig. S7Ba-Bb.

### Assumptions for the CA3 network model

We consider an animal, which experience a series of episodes. We call MF and PP afferent inputs to CA3-PCs carrying the information of *i*-th episode as *i*-th pattern (***P***_i_).

1. *MF inputs are strong and sparse, while PP inputs are weak and diffuse*: Only a small and non-overlapping subset of CA3-PCs receive MF input during encoding (or learning) phase, while almost all CA3-PCs receive PP inputs with different strength. Henceforth, a group of CA3-PCs that receive MF inputs carrying information of the *i*-th pattern (***P*_i_**) will be denoted as **M_i_** subset.
2. *Distributions of MF and PP input strength are skewed*: During encoding phase, the MF input strength to CA3-PCs of a given **M** subset follows a log-normal distribution (Buzsaki & Mizuseki, 2014). We assumed that a small fraction of CA3-PCs in a given **M** subset receive strong MF inputs, in which LTP-IE is induced by concerted actions of intracellular Ca^2+^ and Zn^2+^ signaling (Eom et al., 2019). A subset of CA3-PCs that underwent LTP-IE during encoding of ***P*_i_** will be denoted as **M_i_**\*.
3. *LTP at A/C and PP synapses requires converging MF inputs*: Because A/C and PP synapses display Hebbian plasticity (McMahon & Barrionuevo, 2002; Mishra et al., 2016; Tsukamoto et al., 2003)), LTP at A/C and PP synapses is critically dependent on the postsynaptic firing. Sparse and strong MF input provides an additional variance in the excitatory input distribution over the CA3-PC population and thus governs the firing pattern of CA3-PCs during a learning phase under competitive *k*-Winners-Take-All regime (O’Reilly & McClelland, 1994; Treves & Rolls, 1992). Moreover, acetylcholine (Ach) level is high during encoding phase, and the high Ach reduces the A/C synaptic strength by 50% (Hasselmo, 2006; Vogt & Regehr, 2001). As a result, LTP at A/C and PP synapses is supervised by MF inputs that carry information of an episode. The memory for *i*-th episode (***E*_i_**) is stored at PP and A/C synapses of CA3-PCs comprising **M**_i_ subset (referred to as **M_i_** PCs).

### Data analysis

Electrophysiological data were acquired and digitized using an analogue-to-digital converter (Digidata 1440A, Molecular Devices) and stored in a computer hard disk. Analyses of data were performed with Igor Pro (version 6.10A; WaveMetrics, Lake Oswego, OR, USA) and Clampfit 10.2 (Molecular Devices, Sunnyvale, CA, USA). We used the bias-corrected version of Akaike information criterion (AIC) to select a better distribution model for the histogram data of fluorescence intensities in Fig. S4D (Hurvich & Tsai, 1989). Statistical data are expressed as mean ± standard error of the mean (SEM), and the number of cells measured (denoted as *n*). Statistical data were evaluated for normality and variance equality with Kolmogorov-Smirnov test and Levene’s test, respectively. For data that satisfy normality and equality of variances, statistical evaluations were performed with Student’s t-test or general linear model (GLM). For data that did not satisfied above properties, nonparametric tests, such as Mann-Whitney U test or Wilcoxon signed rank test, were performed to evaluation. For simplicity of description, significance level (P) was marked as P < 0.05. n.s., no statistical significance; ^∗^P < 0.05; ^∗∗^P < 0.01; ^∗∗∗^P < 0.005 in figures. Statistics processing was performed with PASW Statistics 18 (SPSS Inc, 2009, Chicago, IL, USA).

## Results

### 1. Differential effects of intracellular glutathione on somatic and MF-induced LTP-IE

The LTP-IE is mediated by Ca^2+^ and protein tyrosine kinase (PTK)-dependent downregulation of Kv1.2, which is antagonized by receptor protein tyrosine phosphatase alpha (RPTPα) (Fig. S1) (Eom et al., 2019). RPTPα is activated by PKC, and inactivated by oxidative stress and Zn^2+^ (Blanchetot et al., 2002; Meng et al., 2002). Reducing molecules such as reduced form of glutathione (GSH) are normally present in cytosol in millimolar range (Forman et al., 2009). Intracellular perfusion with the conventional whole-cell patch pipette solution lacking reducing molecules may bias the cellular redox to oxidized state. To prevent inactivation of protein tyrosine phosphatase (PTP) during whole-cell recording (WCR), we added 5 mM GSH to the pipette solution. We confirmed that input conductance (G_in_), PP-evoked EPSPs (PP-EPSPs) and action potential (AP) parameters were stable up to 45 min of WCR in the presence of 5 mM GSH (Fig. S2). Henceforth, all the whole-cell patch experiments shown in Fig. 1-4 were done using pipette solution containing GSH, unless stated otherwise.

The long lasting increases in input resistance (inverse of G_in_) and PP-EPSPs are the hallmarks of LTP-IE in CA3-PCs, which can be induced by 10 Hz somatic AP firing (somatic conditioning, SC) or 20 Hz MF stimulation (MF conditioning)(Eom et al., 2019; Hyun et al., 2015; Hyun et al., 2013). After measuring the baseline G_in_ for 5 min, the somatic or MF conditioning was delivered (black or red arrowhead, respectively). To our surprise, in the presence of GSH, the SC-induced decrease in G_in_ was not sustained (Fig. 1Aa-c), whereas it was without GSH (Hyun et al., 2013). This GSH effect was not observed after MF conditioning. The acceleration of the 1^st^ AP latency measured from voltage responses to a ramp current (250 pA/s) was in parallel with those of G_in_ (Fig. 1Ad-Ag). Both somatic and MF conditioning increased PP-EPSPs, but faster as previously observed (Hyun et al., 2015) (Fig. 1Ba-Bc). As expected, the SC-induced increase in PP-EPSP was not sustained, whereas those after MF conditioning stayed enhanced (Fig. 1Ba-Bc). Neither somatic nor MF conditioning altered the amplitude of PP-EPSCs (Fig. 1Bd-Be). EPSC amplitude depends on neurotransmitter release and AMPA receptor expression but little on intrinsic excitability. Thus, these results suggest that MF inputs augment CA3 dendritic excitability thereby increasing PP-EPSPs. Given that PTP activity is preserved by GSH in the pipette solution, our results suggest that PTP may rapidly reverse the LTP-IE induced by SC but not that by MF conditioning. This results can be understood in light of our previous findings that MF conditioning induces not only Ca^2+^ but also Zn^2+^ signaling, and the latter facilitates LTP-IE by reversible inhibition of PTP (Eom et al., 2019). Somatic conditioning which induces Ca^2+^ transients (CaTs) without Zn^2+^ signaling, however, may activate both PTKs and PTPs (Brandt et al., 2003; Eom et al., 2019; Lev et al., 1995; Tsai et al., 1998), and lead to the short-lasting potentiation of intrinsic excitability (PIE). This result is in stark contrast to our previous observation that SC readily induced long-lasting PIE in the absence of GSH (Hyun et al., 2013).

### 2. Depotentiation of MF-induced potentiation of PP-fEPSPs by A/C synaptic stimulation

We hypothesized that CaTs without Zn^2+^ signaling may reverse MF-induced LTP-IE when the activity of PTP is preserved. CaTs without Zn^2+^ signaling can be evoked by stimulation of soma or afferent fibers other than MFs (non-MF) such as PP or associational/commissural (A/C) fibers (Eom et al., 2019). To prevent perturbation of the intracellular milieu of CA3-PCs, we monitored PP-evoked field EPSPs (PP-fEPSPs) before and after MF conditioning and subsequent A/C fiber stimulations. The recording and stimulating electrodes were placed as shown in Fig. 2Aa (see Materials and Methods). Stimulation of PP evoked a negatively deflecting fEPSP at the recording electrode placed in stratum lacunosum-moleculare (SLM) (gray-black, Fig. 2Ab), while stimulation of MFs (cyan) and A/C fibers (red) evoked positive fEPSP at SLM. To evoke antidromic firing of CA3-PCs and activate A/C synapses, the alveus of CA3 was stimulated with 40 pulses for 2 s, which induced AP bursts following EPSP summation in the CA3-PC under WCR (Fig. 2B). Both EPSPs and AP bursts were abolished by bath application of CNQX, suggesting that antidromic firings of CA3-PCs activated A/C inputs to the CA3-PC under WCR, resulting in AP bursts.

We regarded the maximal slope over an initial half of rising phase of fEPSP as an initial slope of PP-fEPSP. The initial slope of fEPSP represents fast EPSC, while the peak fEPSP amplitude reflects delayed activation of somato-dendritic currents as well as synaptic current (Garcia et al., 2010). Fig. 2Ca-b shows the input-output relationships for the peak amplitude and initial slope of PP-fEPSP, respectively. The latency of PP-fEPSPs was not altered by the increase in stimulation intensity (Fig. 2Cc). The PP-fEPSPs were attenuated by bath application of DCG-IV, an agonist of group II mGluR, which is highly expressed in PP (Shigemoto et al., 1997)(Fig. 2Cd).

MF conditioning enhanced the peak amplitude of PP-fEPSP, but not the initial slope of PP-fEPSPs (Fig. 2Da and 2Ea), consistent with our patch clamp findings that MF conditioning enhanced PP-EPSPs, but not PP-EPSCs (Fig. 1). Indeed, the alveus stimulation (10 pulses at 50 Hz, A/C conditioning) applied at 20 min after MF conditioning induced gradual decrease in the peak amplitude of PP-fEPSPs (Fig. 2D). The initial slope of PP-fEPSPs and paired pulse ratio (PPR) of slopes were not affected by MF and A/C conditioning (Fig. 2E), suggesting that changes in dendritic excitability of CA3-PCs underlie the MF-induced potentiation and A/C-induced depotentiation of PP-fEPSPs.

### 3. MF-induced LTP-IE is de-potentiated by somatic or A/C synaptic stimulation, and is hindered by Zn^2+^

To test whether the results of field recordings can be replicated in whole-cell recordings, we examined the effects of alveus stimulation on MF-induced LTP-IE. The MF conditioning decreased the G_in_ of CA3-PCs (73.5 ± 1.8%, n = 21, Fig. 3Aa-Ad). The G_in_ of CA3-PCs was restored to the baseline by a train of stimuli (10 pulses at 20 Hz) delivered to the alveus at 15 min after MF conditioning (Fig. 3Aa), consistent with the results of field recordings (Fig. 2). To avoid inadvertent recruitment of various neuromodulator pathways by alveus stimulation, we tested whether direct somatic stimulation can depotentiate MF-induced LTP-IE. Somatic AP bursts comprised of 10 spikes at 100 Hz, a typical burst firings of CA3-PCs in vivo (Kowalski et al., 2016), applied at 15 min after MF conditioning also restored the G_in_ to the baseline (Fig. 3Ab). To assess the minimal somatic stimulation required for depotentiation, we examined the effects of somatic stimulation with fewer number of spikes. Surprisingly, the bursts of two APs at 100 Hz restored G_in_ to the baseline level (Fig. 3Ac). We examined depotentiation of PP-EPSPs too. Similar to the G_in_ changes, MF-induced potentiation of PP-EPSPs (197.5 ± 19.9%, n = 21, Fig. 3Ba-Bd) was reversed by AP bursts elicited by alveus stimulation (Fig. 3Ba) or by direct injection of ten (Fig. 3Bb) or two (Fig. 3Bc) supra-threshold somatic current pulses at 100 Hz. The amplitudes of PP-EPSCs were not altered after MF conditioning or depotentiation stimulation (time, F_(2,18)_ = 0.017, p = 0.897; stimuli, F_(2,18)_ = 0.718, p = 0.501; stimuli × time, F_(2,18)_ = 0.299, p = 0.745; GLM and simple effect analysis). The changes of G_in_ and PP-EPSPs before and after MF conditioning and depotentiation stimulation are summarized in Fig. 3Ad and Bd, respectively. Somatic AP bursts did not depotentiate MF-induced LTP-IE in the presence of 100 nM ZnCl_2_, which was added to aCSF 10 min after MF conditioning (Fig. 3C), indicating that lack of Zn^2+^ signaling during AP bursts is essential for depotentiation. This result supports our hypothesis that CaTs without Zn^2+^ signaling activates PTP and triggers depotentiation of LTP-IE.

### 4. MF-induced LTP-IE is reversed by homosynaptic induction of LTP at PP-CA3 synapses

MF-induced LTP-IE heterosynaptically potentiates PP-EPSPs, but not PP-EPSCs, because LTP-IE enhances dendritic excitability with little effect on synaptic transmission *per se*. We will refer to homosynaptic LTP of PP-EPSCs as PP-LTP, distinct from MF-induced hetero-synaptic LTP of PP-EPSPs accompanied with LTP-IE. Previously we showed that LTP-IE lowers the threshold for PP-LTP in the mice hippocampal CA3 (Eom et al., 2022). We confirmed this notion in the rat CA3 (Fig. S3 and Supplementary Results).

Above results suggest that MF-induced LTP-IE can be reversed by somatic burst firings elicited by somatic or A/C stimulation (Fig. 2-3), and that MF-induced LTP-IE enables weak PP inputs to elicit APs and induce Hebbian PP-LTP (Fig. S3). We posed a question whether AP firing elicited by weak PP inputs in a MF-conditioned CA3-PC can reverse LTP-IE in the presence of GSH. To this end, we simultaneously monitored PP-EPSPs (Fig. 4Aa) and G_in_ (Fig. 4Ba), while MF conditioning and subsequently weak high-frequency stimulation of PP (PP-HFS) were delivered to induce LTP-IE and PP-LTP, respectively. MF conditioning (Fig. 4Ab) enhanced PP-EPSPs but not PP-EPSCs (Fig. 4Aa), and decreased G_in_ (Fig. 4Ba). At 5 min after MF conditioning, we delivered PP-HFS, which readily evoked AP firings (Fig. 4Ac) unlike naïve CA3-PCs (Fig. S3Aa). After PP-HFS, both the PP-EPSPs (*left*) and PP-EPSCs (*right*) were increased (Fig. 4Ca). However, the G_in_ that had been reduced by MF conditioning was restored to the baseline level (Fig. 4B). The EPSP/EPSC ratio that had been increased by MF conditioning returned to the baseline after PP-HFS as shown in Fig. 4Cb. These results indicate that high-frequency MF inputs prime CA3-PCs for PP-LTP expression by lowering the LTP threshold so that subsequently incoming high-frequency PP inputs readily elicit AP bursts in the primed PCs, and that the PP-induced AP bursts lead to both PP-LTP induction and termination of MF-induced LTP-IE.

### 5. LTP-IE occurs in hippocampal CA3 region, and lasts less than 3 hours in vivo

We examined whether LTP-IE occurs in CA3 ensemble cells activated when rats are exposed to a novel context. We labeled an ensemble of CA3-PCs activated by contextual fear conditioning (CFC) using AAV-RAM-d2tTA-TRE-EGFP (Sorensen et al., 2016). Rats were kept on doxycycline-free diet for 2 days before being subject to CFC. Just after CFC, rats were injected with doxycycline (10mg/kg of body weight), and kept in their home-cage for 1, 1.5 or 3 hours before *ex vivo* examination of G_in_ in CA3-PCs (Fig. 5Aa). At about 30 min WCR of EGFP(+) CA3-PCs, we could observe the diffusion of EGFP into the patch pipette (Fig. 5Ab-d). Fig. 5B shows the baseline G_in_ as a function of EGFP fluorescence intensity. Most EGFP(-) cells (40 out of 45 cells) showed high baseline G_in_ [denoted as EGFP(-/high), black symbols], but a few of them (5 cells) showed low G_in_ [EGFP(-/low), yellow symbols]. The average baseline G_in_ of EGFP(+) CA3-PCs at 1 hour [red symbols, denoted as EGFP(+/1hr) cells] and at 1.5 hours [blue symbols, EGFP(+/1.5hr) cells] were significantly lower than those of EGFP(-/high) cells, and not different from G_in_ of GFP(-/low) cells. On the other hand, G_in_ of EGFP(+) cells at 3 hours [green symbols, GFP(+/3hr) cells] were not different from those of GFP(-/high) cells. Therefore, cells were categorized into high and low G_in_ cell groups: GFP(-/high) and GFP(+/3hr) cells *vs*. GFP(-/low), GFP(+/1hr) and GFP(+/1.5hr) cells under the significance level of 0.05 [F_(4,,72)_ = 38.667, p < 0.001, Tukey HSD]. Since most ensemble cells displayed lower G_in_, the GFP(-/low) cells may represent CA3 ensemble cells that are not infected by AAV.

The inducibility of LTP-IE was evaluated by comparing G_in_ at 30 min after somatic conditioning (denoted as post-SC G_in_) with baseline values. in the low G_in_ cell group, SC did not induce additional decrease of G_in_ (Fig. 5C), while LTP-IE was readily induced in the high G_in_ cell group [baseline vs. post-SC, p < 0.001 in GFP(-);p < 0.554 in GFP(+/3hr); GLM and simple effect analysis]. Moreover, the post-SC G_in_ values were not different among all groups (baseline, F_(3,24)_ = 14.616, p < 0.001; post-SC, F_(3,24)_ = 0.706, p = 0.558, GLM and univariate analysis), suggesting that GFP(-/low), GFP(+/1hr) and GFP(+/1.5hr) cells may have already undergone LTP-IE. The similarity of GFP(+/3hr) to the GFP(-/high) cells indicates that LTP-IE of CA3-PCs does not last longer than 3 hours *in vivo*, and implies that there may be a mechanism that reverses the increased intrinsic excitability while the animal is kept in their homecage (see Discussion).

### 6. LTP-IE occurs in CA3 ensemble cells and is reversed in a subset of putative twice-activated ensemble cells

Given that the LTP-IE of CA3-PCs is maintained at least for 90 min after CFC, we wondered whether the LTP-IE can be reversed earlier by formation of an additional ensemble in the CA3 area. To ensure labeling of all ensemble cells, we further studied the excitability of ensemble cells using B6.Cg-Tg(Fos-tTA,Fos-EGFP*)1Mmay/J mice (referred to as cfos-shEGFP mice), which express short half-life green fluorescent protein (shEGFP, t_1/2_ = 2 hours) under the control of c-*fos* promoter. We compared the fluorescence intensity distribution of CA3 shEGFP(+) cells in cfos-shEGFP mice exposed only once to a context A (ctxA mice) with that in mice exposed to two contexts (A and B; ctxAB mice) sequentially with a 30 min time interval, and found that the latter mice harbor shEGFP(+) cells with distinctly higher fluorescence intensity (Fig. S4 and Supplemental Results).

We examined whether the input resistance of CA3-PCs has any correlation with their epi-fluorescence intensity *ex vivo* from ctxA and ctxAB mice (Fig. 6Aa). For ctxA mice, excitability was examined at two time points: at 60 and 90 min after being kept in their home cage (Fig. 6Aa; ctxA60 and ctx90 mice, respectively). As shown below, shEGFP(+) CA3-PCs in ctxA60 and ctx90 mice [denoted as ctxA60(+) and ctxA90(+) cells, respectively] were not different in fluorescence intensity and excitability parameters. To correct the depth-dependent fluorescence attenuation in epifluorescence microscopy, we added 50 μM Alexa Flour 594 (AF594; Invitrogen) to a patch pipette solution for ratio-metric measurement of shEGFP and AF594 fluorescence intensities. Consistent with the results of Fig. S4, a subset of shEGFP(+) CA3-PCs from the ctxAB mice displayed higher green-to-red fluorescence (G/R) ratio than the maximal G/R ratio of ctxA60(+) or ctxA90(+) cells, and will be referred to as ctxAB(++) cell (Fig. 6Bc-d). The other subset which had G/R ratio similar to ctxA60(+) and ctx90(+) cells, and will be referred to as ctxAB(+) cell (Fig. 6Ba-b). The G/R ratios of all shEGFP(+) cells could be categorized into two groups: a group comprised of ctxA60(+), ctxA90(+) and ctxAB(+) cells and the other group of ctxAB(++) cells [F_(3,43)_ = 39.741, p < 0.001, ANOVA and Tukey HSD, Fig. 6Be]. Because ctxAB(++) cells were observed only in the mice which visited context A and B, the high G/R ratio of ctxAB(++) cells may be caused by double activation of c-*fos* promoter in the contexts A and B.

The baseline G_in_ values for ctxA60(+) and ctxA90(+) cells were significantly lower than those of GFP(-) cells (p < 0.001 for both), and were not further decreased by SC (ctxA60(+), p = 0.814; ctxA90, p = 0.126), suggesting that these cells have already undergone LTP-IE while the animal visited context A. The baseline G_in_ and LTP-IE inducibility of ctxAB(+) CA3-PCs were similar to those of ctxA60(+) or ctxA90(+) cells [For baseline G_in_, p = 0.819; For pre- vs post-SC G_in_ of ctxAB(+) cells, p = 0.873; Fig. 6Cd-Cf]. In contrast, the excitability of ctxAB(++) CA3-PCs was heterogenous. To test the inducibility of LTP-IE for ctxAB(++) cells, the ratio of post-SC G_in_ over baseline G_in_ was measured. The ctxAB(++) cells were plotted on the plane of the G_in_ ratio *vs.* baseline G_in_, and were clustered using *k*-means algorithm, yielding two clusters shown in Fig. 6Cc. The low and high baseline G_in_ groups are referred to as ctxAB(++/1) cells (n = 11) and ctxAB(++/2) cells (n = 9), respectively (Baseline G_in_, t = 4.775, p < 0.001; G_in_ ratio, t = −8.115, p < 0.001). The baseline G_in_ and the G_in_ ratio are shown as a function of G/R ratio in Fig. 6Cd and in Fig. 6Ce, respectively. Both baseline G_in_ and G_in_ ratio of shEGFP(+) cells were categorized into two groups: GFP(-) and ctxAB(++/2) *vs*. ctxA60(+), ctxA90(+), ctxAB(+) and ctxAB(++/1), which will be referred to as low and high excitability groups, respectively [For baseline G_in_, F_(5,32)_ = 12.883, p < 0.001; For G_in_ ratio, F_(5,32)_ = 19.565, p < 0.001; ANOVA and Tukey HSD]. The mean G_in_ values before and after SC are summarized in Fig. 6Cf. The baseline G_in_ values within high excitability groups were not different one another, but significantly lower than the baseline G_in_ of GFP(-) cells [baseline G_in_ (gray bars), F_(5,31)_ = 13.053, p < 0.001; post-SC G_in_ (red bars), F_(5,31)_ = 1.213, p = 0.364, Univariate analysis], and rather similar to the post-SC G_in_ of GFP(-) cells (χ^2^ = 7.279, df = 3, p = 0.064, Kruskal-Wallis test).

The 1st AP latency and PP-EPSP/EPSC ratio reflect the excitability of soma and distal apical dendrite, respectively. Comparing these parameters in different cell groups, we found that both of them are highly correlated with the baseline G_in_: faster 1st AP latency and high EPSP/EPSC ratio in high excitability cell group and vice versa (Fig. S5 and S6, respectively; see Supplemental Results). These results suggest that CA3 ensemble cells except ctxAB(++/2) cells had already underwent LTP-IE *in vivo*. In contrast, ctxAB(++/2) cells were not different from GFP(-) cells with regard to all aspects: baseline G_in_, the 1^st^ AP latency, and EPSP/EPSC ratio, and their pre- vs post-SC ratios (Fig. 6Cc, S5Ba and S6E), suggesting that ctxAB(++/2) cells are not different from GFP(-) cells in somatic and distal dendritic excitability, even though they are supposed to be recruited to ensembles A and B twice. We discussed below possible mechanisms underlying the heterogeneity of ctxAB(++) cells (see Discussion).

### 7. Lack of LTP-IE in the ensemble CA3-PCs of ZnT3 hetero-knockout mice

The results of the present study together with Eom et al. (2019) indicate that MF-induced Zn^2+^ signalling inhibits PTP, and thus widens the cytosolic [Ca^2+^] window for LTP-IE induction and maintains LTP-IE in the normal redox state. We tested whether LTP-IE occurs *in vivo* when MF-induced Zn^2+^ signalling is disrupted by hetero-insufficiency of ZnT3, a vesicular Zn^2+^ transporter. To this end, ZnT3 hetero-knockout(HT) x cfos-shEGFP hybrid mice (ZnT3HT/cfos-EGFP) were made by crossing ZnT3-knockout (KO) mice with cfos-shEGFP mice. We confirmed that LTP-IE was induced by 20 Hz MF stimulation but not by 50 Hz stimulation in CA3-PCs of ZnT3HT mice (n =5, t = −1.141, p = 0.318, paired t-test; Fig. 7A), similar to those of ZnT3-KO mice (Eom et al, 2019). Next, we examined correlation of G_in_ with G/R ratio in ZnT3HT/cfos-shEGFP mice which experienced context A and B (Fig. 7B). Unlike wildtype CA3-PCs, CA3-PCs of ZnT3HT/cfos-shEGFP mice exhibited high G_in_ regardless G/R ratio (Fig. 7C and Da). Moreover, 10 Hz somatic conditioning induced LTP-IE in high G/R ratio cells as well as GFP(-) cells (Fig. 7C and Db), indicating that induction and/or maintenance of LTP-IE are impaired in vivo in ZnT3-HT mice. Given that dendritic Zn^2+^ transients in CA3-PCs were induced by MF inputs but not by PP or A/C inputs, these results suggest that CA3-PCs that received high-frequency MF inputs undergo LTP-IE in vivo.

### 8. Computational modelling of the association of two sequential events in the CA3 network

Our results suggest that high-frequency MF inputs prime a CA3-PC, and high-frequency PP inputs to a primed CA3-PC not only induce homosynaptic LTP and but also reverse the prior MF input-induced LTP-IE (Fig. 4). To conceive possible roles of this learning rule in the CA3 network dynamics, we incorporated it into a CA3 recurrent network model. The assumptions adopted for the network model is based on the Hebb-Marr network (McNaughton & Morris, 1987) and the computational theory of CA3 function (Lee & Kesner, 2004; Treves & Rolls, 1992), and are described in detail in *Materials and Methods*.

To implement the LTP-IE-based learning rules, we constructed a computational model of recurrent network comprised of 1000 PCs and 200 interneurons (INs) *in silico*, which are numbered with indices from 1 to 1000 and 1001 and 1200, respectively. Fig. 8A and Fig. S7A show the schematic synaptic connections between PCs and INs, and the initial connectivity map of the model network, respectively. We assumed that PCs make excitatory synapses with each other with the probability of 10%. The synaptic weights between PCs were randomly set to a value between 0 and 0.35 nS. The spike properties of PCs and INs were assumed to follow a leaky integrate and fire model using parameter sets as described in Materials and Methods.

We considered an animal which experiences a series of episodes. During each episode, MF and PP afferent inputs are assumed to present the information of *i*-th episode as *i*-th input pattern (denoted as ***P***_i_) to CA3-PCs. The model network was trained by four patterns that arrive at the network in a sequence with different time intervals (0.8-1.2 s) as shown in Fig. 8Ba (We adopted here short time intervals to lessen computational time, but longer time intervals do not alter the main conclusion). Each input pattern is comprised of sparse non-overlapping MF and diffuse PP inputs (red and blue dots in Fig. 8Ba, respectively). The arrival of MF inputs in each pattern is confined in a subset of 200 PCs, which constitute an **M** subset, while PP inputs arrive over all the PCs in the network. We assumed that each PC receives different numbers of MF or PP inputs per a pattern, which are randomly given under a log-normal distribution (Ikegaya et al., 2013) (Fig. 8Ba and Fig. S7D). During the encoding phase, PP and A/C synapses of PCs undergo plastic changes under the spike timing dependent plasticity (STDP) rules. We assumed symmetric and asymmetric STDP curves for A/C and PP synapses, respectively (Mishra et al., 2016) (Fig. S7B).

Our results suggest that coincident increases in cytosolic [Ca^2+^] and [Zn^2+^] induce LTP-IE, while [Ca^2+^] increase alone reverses LTP-IE. In our simulation, the [Ca^2+^] and [Zn^2+^] thresholds required for triggering LTP-IE were set such that MF input-induced Ca^2+^ and Zn^2+^ transients cross the threshold in about 25% of PCs of an **M** subset during encoding phase. A subset of CA3-PCs that undergo LTP-IE during encoding of ***P_i_*** will be denoted as **M_i_\***. Exemplar traces for [Ca^2+^]_i_ and [Zn^2+^]_i_ simulated in an **M_1_\*** PC are shown in Fig. 8C. To simulate the time course of excitability change we defined an intrinsic excitability parameter, *IE*, which varies from 0 to 1. *IE* is equal to 0 when a cell is in a resting state, and it starts to increase toward one when both [Ca^2+^]_i_ and [Zn^2+^]_i_ co-incidentally surpass the Ca^2+^ and Zn^2+^ thresholds (Caθ1 and Znθ, respectively). Once *IE* is increased by MF inputs, it stays high until a CaT that surpasses another Ca^2+^ threshold (Caθ2) occurs devoid of a ZnT. PP inputs to such PCs with high *IE* can readily induce APs independent of MF inputs. To simulate this, a transient current was injected upon arrival of PP inputs to a primed PC that has *IE* > 0.7 (Fig. S7C)(Jahnke et al., 2015). The PP input to a primed cell readily elicits APs in a postsynaptic PC resulting in a CaT devoid of Zn^2+^ signaling. When the peak of a PP-induced CaT is higher than a given Ca^2+^ threshold (Caθ2), the *IE* returns to zero (de-potentiation; Fig. 8C). Fig. 8Bb shows the simulation results for the firing of PCs in response to the four sequentially incoming input patterns shown in Fig. 8Ba. The PP inputs of ***P*_2_** readily elicit firings of the **M_1_\*** PCs (subset of primed PCs in **M**_1_), and thus fire together with PCs of the **M_2_** subset. These concomitant firing of PCs in **M_1_\*** and **M_2_** subsets allows recurrent synapses to be strengthened between the two orthogonal ensembles. Fig. 8D shows a matrix of recurrent synaptic weights consequent to these auto- and hetero-associative learning rules.

Next, we tested whether the recurrent network trained under these learning rules recalls the whole sequence of memories from a partial cue of ***P***_1_. We assumed no reduction in A/C synaptic weight during recall, because recall occurs under the low cholinergic state. To trigger pattern completion-based recall of the memory of ***P***_1_, which is represented by **M_1_**, a burst of PP inputs carrying a part of ***P*_1_** (denoted as ***p*_1_**) was applied to the network (Fig. S7D). The synaptic weights of the PP inputs of ***p_1_*** have been already strengthened during encoding phase only in **M_1_** PCs. The PP inputs of ***p*_1_** drive firing in a few **M_1_** PCs (14 PCs in the example shown in Fig. 8E), which propagated through already strengthened recurrent connections, resulting in pattern completion-based recall of **M_1_** (Fig. 8E). Meanwhile, the activation of PCs in **M_1_**\* plays a role of partial cue to **M_2_**, and triggers the second pattern completion-based recall of **M_2_**. In this way, the PP inputs carrying a partial cue of ***P***_1_ trigger the recall of whole sequence of memories that were hetero-associated during encoding phase (from **M_1_** to **M_4_**). This simulation implies that LTP-IE may allow the CA3 network to recall of memories of sequential events, even if they are temporally discontiguous. The simulation results are summarized in Supplemental Results.

## Discussion

### LTP-IE may occur in the CA3 ensemble cells

The present study suggests that LTP-IE is de-potentiated not only by PP inputs probably during the encoding phase (Fig. 4) but also without any episodic events within 3 hours (Fig. 5). A/C input strength is critically modulated by acetylcholine (Ach) level. High Ach level in the hippocampus during awake and active learning state enhances MF synapses and suppresses A/C synapses (Vogt & Regehr, 2001). When Ach level is low during a retrieval or consummatory phase, the suppression of A/C synapses is relieved resulting in spontaneous offline reactivation of neuronal ensembles, manifested as sharp wave ripples (SWRs). Unlike PP-EPSPs, A/C-EPSPs are not enhanced by LTP-IE (Hyun et al., 2015). In the low cholinergic state, however, A/C input-driven firings of CA3-PCs would be facilitated (Kowalski et al., 2016; Malezieux et al., 2020), and may contribute to depotentiation of LTP-IE even if A/C inputs do not induce IEG activation. This possibility may be responsible for depotentiation of LTP-IE while animals were kept in the homecage. It remains to be tested whether the A/C synaptic inputs to CA3-PCs during low cholinergic state contribute to de-potentiation of LTP-IE.

Immediate early gene (IEG)-induced expression of fluorescence protein has been widely used for labeling neuronal ensembles to investigate the recall of memory represented by the labeled ensemble. To investigate whether LTP-IE is reversed by the second activation of ensemble cells, it is essential to distinguish once- *vs*. twice-activated ensemble cells in an animal that experienced two contexts within 2 hours because LTP-IE is spontaneously reversed less than 3 hours (Fig. 5). No currently available labeling technique meets these requirements, although techniques for labeling active population of neurons with high temporal precision has been recently developed (Lee et al., 2017; Moeyaert et al., 2018). Thus, we compared the expression of c-fos promoter-induced shEGFP expression level between cfos-shEGFP transgenic mice which were exposed a novel context once and twice (denoted as ctxA and ctxAB mice). Because a subset of ensemble CA3-PCs from the ctxAB mice expressed distinctly high level of shEGFP compared to those from the ctxA mice observed by both confocal and epifluorescence microscopy (Fig. S4 and Fig. 6), we regarded such shEGFP(++) subset as twice-activated ensemble cells. Nevertheless, it cannot be ruled out that it contains once-activated ensemble cells.

### Why a subset of ctxAB(++) cells exhibit G_in_ similar to that of non-ensemble cells?

CA3-PCs with distinctly high EGFP expression from ctxAB mice were comprised of mixed population in the excitability (Fig. 6 and Fig. S5-S6). Why ctxAB(++/2) cells exhibit the normal excitability despite that these cells may participate in both ensembles encoding contexts A and B? There are two possibilities for the normal excitability of ctxAB(++/2) cells. First, these cells may have never experienced LTP-IE. Alternatively, they underwent LTP-IE in context A, and the LTP-IE is de-potentiated in context B. The first possibility is unlikely because most CA3 ensemble cells of the animal that visited only context A [ctxA60(+) and ctx90(+) cells] exhibited high excitability. Previously we showed that MF inputs are privileged for the induction of LTP-IE (Eom et al, 2019). Moreover, Figure 1 suggests that, even if the excitability would be enhanced by non-MF inputs, the high excitability state cannot be sustained longer than 30 min under the physiological redox conditions. This notion is supported by the ex vivo results that the CA3 ensemble cells in ZnT3- heteroKO mice exhibited no sign of LTP-IE (Fig. 7). Since we examined G_in_ at 60 min after the last visit, ctxAB(++) cells as well as ctxA60(+) and ctxA90(+) cells may have received high frequency MF inputs while the animal visited context A, assuming that ctxAB(++) are twice-activated ensemble cells. If the latter possibility is correct, the mechanism underlying de-potentiation of LTP-IE can be explained by our findings that non-MF inputs can reverse the MF-induced high excitability state of CA3-PCs (Fig. 2, 3 and 4). Moreover, CA3-PCs in the primed state readily fire in response to subsequently arriving high frequency PP inputs, which also reverse the high excitability state (Fig. 4). Given that ctxAB(++) cells are activated both in context A and B, their firing in context B can be elicited not only by MF inputs but also by PP inputs alone as long as the LTP-IE is maintained while the animal visited context B. Therefore, ctxAB(++) cells may be comprised of two groups: CA3-PCs that received high frequency MF inputs in both contexts, and those received MF inputs in context A and subsequently only PP inputs in context B. The latter group may undergo MF-induced LTP-IE in context A and PP-induced depotentiation in context B, and thus eventually exhibit normal baseline G_in_, because PP inputs can reverse the prior MF input-induced LTP-IE (Fig. 4).

### Computational implications

Hippocampal CA3 has been implicated not only in non-overlapping representation of episodic memories but also in the memory of sequence of temporally discontiguous events (Farovik et al., 2010). Recently, the synaptic tagging and capture hypothesis and/or post-learning CREB-dependent modification of intrinsic excitability have been proposed as network mechanisms that permit the overlap of ensembles representing temporally separate episodes in hippocampal CA1 and amygdala (Lisman et al., 2018). This overlap may be mediated by increased sharing of feedforward inputs between the ensembles because recurrent connections between principal cells lack in these areas. The same regulation of ensemble overlap in a recurrent network would have a detrimental effect on the orthogonality of ensembles because co-active neurons tend to be strongly connected via recurrent synapses (called auto-association), which may eventually lead to a merge of ensembles. Therefore, different mechanisms may be necessary in order for two orthogonal ensembles to be associated (called hetero-association) in a recurrent network, but it has been little understood. The LTP-IE-based hetero-association of ensembles proposed in the present study prevents not only uncontrolled overlap between non-adjacent ensembles in a sequence (for example, link of **M_1_** and **M_3_**), but also further hetero-association between ensembles carrying information non-adjacent events in a sequence. Therefore, the learning rule may allow two orthogonal ensembles adjacent in a sequence to be linked with the overlapping fraction tightly regulated.

## Acknowledgements

This study was supported by National Research Foundation of Korea (Grant/Award Number to SHL, 2020R1A2C2006438).

## Abbreviations

A/C: associational/commissural
Ach: acetylcholine
CA3-PC: CA3 pyramidal cell
CaTs: Ca^2+^ transients
CFC: contextual fear conditioning
CREB: cAMP response element binding protein
E-S: EPSP-to-spike
fEPSP: field EPSP
G_in_: input conductance
GSH: reduced form of glutathione
KO: knock-out
LTP-IE: long- term potentiation of intrinsic excitability
MF: mossy fiber
PIE: potentiation of intrinsic excitability
PP: perforant pathway
PTK: protein tyrosine kinase
PTP: protein tyrosine phosphatase
SC: somatic conditioning
shEGFP: short-half life EGFP
WCR: whole-cell recording

