## supplementary materials for "Input-specific bi-directional regulation of CA3 pyramidal cell excitability: its implications in sequence learning"

### Supplemental Results

#### *Determination of PP-LTP threshold before and after somatic or MF conditioning (Fig. S3)*

We examined the role of LTP-IE in metaplastic regulation of PP inputs using GSH-free pipette solution (Fig. S3). To this end, we first determined the threshold of stimulation strength for PP-LTP induction adopting the same HFS protocol as (McMahon & Barrionuevo, 2002), which consists of 10 bursts repeated every 10 s with each burst being composed of 20 pulses at 100 Hz (PP-HFS, Fig. S3Aa and S4Ba). Testing PP stimulation of different intensities, we found that PP-LTP requires strong PP inputs sufficient to elicit somatic APs during the LTP induction (Fig. S3A and S4B). Because PP-EPSPs are affected by dendritic excitability, PP stimulation intensity was quantified as the baseline PP-EPSC amplitude. The potentiation ratio was measured as the baseline-normalized PP-EPSC amplitudes 30 min after PP-HFS, and plotted as a function of corresponding PP stimulation intensity (quantified as baseline EPSC amplitudes) in Fig. S3C. We fitted a logistic function to this relationship. We regarded more than 50% increase of PP-EPSC amplitude at 30 min of PP-HFS compared to the baseline value as successful PP-LTP induction (Chen et al., 2006). The x-value of the fitted function at 150 was read as 12.55 pA. The PP-HFS with baseline EPSC < 12.55 pA did not induce PP-LTP (Fig. S3Ab), whereas PP-HFS with EPSC > 12.55 pA did (Fig. S3Bb). Next, we tested whether weak PP-HFS (baseline PP-EPSC < 12.55 pA) which does not induce PP-LTP in a naïve PC, can trigger PP-LTP in the CA3-PCs that have undergone MF or somatic conditioning. Despite of the weak stimulation intensity, PP-HFS readily elicited APs in the CA3-PCs, which underwent MF or somatic conditioning, indicative of E-S potentiation (Fig. S3Da, *upper* and *lower*, respectively). Consistent with (Hyun et al., 2015), neither MF conditioning nor somatic conditioning (arrowhead, Fig. S3Db) altered PP-EPSCs (Fig. S3Dc, Ea, Eb), but both of them enhanced PP-EPSPs that were measured 6 min after the conditioning (Fig. S3Dc, Ec, Ed). Distinct from the naïve CA3-PCs (Fig. S3Aa), weak PP-HFS readily induced PP-LTP in conditioned CA3-PCs (Fig. S3Db, Ea, Eb). The PP-EPSPs in the conditioned CA3-PCs were also enhanced after the weak PP-HFS (Fig. S3Ec, Ed). For the conditioned CA3-PCs, the potentiation ratios are plotted as a function of baseline EPSC amplitudes in Fig. S3C, showing that the threshold for PP-LTP induction is lowered in the CA3-PCs compared to naïve CA3-PCs.

*Higher GFP expression in twice-activated ensemble CA3-PCs of cfos-shEGFP mice (Fig. S4)*

We hypothesized that, when a neuron is twice activated by two sequential events with a given time interval, the neuron would have higher fluorescence intensity than other neurons activated by a single event. To test this hypothesis, we examined distribution of fluorescence intensities of CA3-PCs from cfos-shEGFP mice which visited only context A and from those which visited two contexts A and B sequentially in a 30 min interval (referred to as ctxA and ctxAB mice, respectively; Fig. S4). Mice visited each context for 5 min, and otherwise were kept in homecage. At 90 min after the first visit, they were sacrificed for preparation of brain slices. We examined fluorescence intensity of shEGFP(+) CA3-PCs located at a depth less than 60 μm from the surface of hippocampal slices. Fig. S4B shows representative fluorescence intensity profiles of z-stack confocal images of shEGFP(+) CA3-PCs from ctxA mice (72 cells, Fig. S4Ba-b) or from ctxAB mice (77 cells, Fig. S4Bc-Bd). The scatter plot of peak fluorescence intensity of each shEGFP(+) CA3-PCs as a function of its depth in the slice revealed that the CA3 area of ctxAB mice harbours CA3-PCs displaying distinctly higher fluorescence intensity compared to that of ctxA mice (Fig. S4C). The distribution of fluorescence intensities of shEGFP(+) CA3-PCs in either group of mice showed negative skewness (Fig. S4D), indicative of log-normal (LN) distribution. The fluorescence intensity distribution of shEGFP(+) cells of ctxAB mice, but not that of ctxA mice, was better fitted by bimodal LN distribution (Fig. S4D; AIC for mono- vs. bi-LN fits = 24.8 *vs*. 26.6 for ctxA; 2.24 *vs*. -9.12 in ctxAB; Note that a model with lower AIC is preferred), suggesting that shEGFP(+) cells of high intensity group are unique to the CA3 ensemble cells of ctxAB mice.

*The baseline AP onset time and PP-EPSP/PP-EPSC ratio are highly correlated with the baseline input conductance (Fig. S5 and Fig. S6)*

The AP onset time upon injection of a short current pulse (400 - 600 pA, 8 - 12 ms; Fig. S5a and S5b) and the latency of first AP evoked by triangular ramp current (250pA/s for 1 s; Fig. S5Ba and S5Bb) were measured before and after somatic conditioning (SC). Compared to GFP(-) cells, the cells of high excitability group exhibited not only lower $G_{in}$ but also faster AP onset time and the first AP latency (Fig. S5A), reflecting downregulation of D-type K current (Hyun et al, 2013). These two AP parameters measured in the cells of high excitability group were not different from those in GFP(-) cells that underwent LTP-IE (AP onset, $\chi^2 = 1.195$, df = 3, p = 0.754; 1$^{st}$ AP latency, $\chi^2 = 3.149$, df = 3, p = 0.369, Kruskal-Wallis test), suggesting that the CA3 ensemble cells had already underwent LTP-IE *in vivo*. In contrast, ctxAB(++/2) cells were not different from GFP(-) cells with regard to AP onset time and AP latency before and after somatic conditioning (baseline AP onset time,  p =0.419;  AP latency, p = 1.00).

Next, We examined PP-EPSPs in CA3-PCs *ex vivo* from cfos-shEGFP mice that have experienced context A or A/B as shown in Fig. 6A. Because LTP-IE enhances the excitability of distal apical dendrite rather than PP-EPSC (Hyun et al., 2015), we compared the EPSP/EPSC ratio between different cell groups. To this end, the stimulation intensity was adjusted such that PP-EPSC amplitudes are largely in the range of 10 to 20 pA and corresponding PP-EPSPs were measured in the same cell. Similar to the distribution of $G_{in}$ as a function of G/R ratio (Fig. 6Cd), the EPSP/EPSC ratio was higher in ctxA60(+), ctx90(+), and ctxAB(+) cells than GFP(-) cells, and those of ctxAB(++) cells were heterogenous (Fig. S6Aa). The plot of the baseline EPSP/EPSC ratio as a function of baseline $G_{in}$ revealed highly negative correlation of the baseline EPSP/EPSC ratio to the baseline $G_{in}$ (Pearson's r = 0.795, p < 0.001, Pearson correlation test; Fig. S6Ab). Similar to the $G_{in}$ ratio of ctxAB(++) cells (Fig. 6Cc), the EPSP/EPSC ratios of those cells were split into two clusters. Because ctxAB(++) cells with low $G_{in}$ exhibited high EPSP/EPSC ratio and vice versa (Fig. S6Ab), ctxAB(++) cells exhibiting high EPSP/EPSC ratio were regarded as ctxAB(++/1) cell, and the other subset as ctxAB(++/2) cell. Similar to $G_{in}$, EPSP/EPSC ratio of shEGFP(+) cells were categorized two groups: low and high excitability groups, which are comprised of GFP(-) and ctxAB(++/2) cells *vs*. ctxA60(+), ctxA90(+), ctxAB(+) and ctxAB(++/1) cells, respectively ($F_{(5,33)} = 24.864$, p < 0.001; one-way ANOVA and Tukey HSD). Fig. S6B shows representative traces for PP-EPSC and PP-EPSP, and the time course for the EPSP/EPSC ratio changes before and after somatic conditioning in five cell groups (the same colour codes as in *A*). Both baseline and post-SC PP-EPSC amplitudes were not different between cell groups (Fig. S6Ba and Ca). However, the baseline PP-EPSP amplitudes were significantly higher in the high excitability group than the low excitability group [low *vs*. high excitability group, $F_{(5,33)} = 24.864$, $p < 0.001$; 1-way ANOVA and Tukey HSD; Fig. S6Bb and Cb]. For the cells of high excitability group, somatic conditioning did not further enhance the PP-EPSP while it did in the cells of low excitability group (Fig. S6Bc and Cb). The baseline EPSP/EPSC ratios of cells in the high excitability group were not different from post-SC EPSP/EPSC ratio measured in GFP(-) and ctxAB(++/2) cells ($\chi^2 = 1.365$, df = 3, p = 0.714, Kruskal-Wallis test; Fig. S6Bc]. These notions are also evident from the plot of individual cells on the plane of PP-EPSP vs. PP-EPSC amplitude in Fig. S6D. The baseline EPSP/EPSC ratio was closely correlated with inducibility of LTP-IE probed by potentiation of EPSP/EPSC ratio (Fig. S6E). In this plot, cells were also categorized into two groups by *k*-means clustering. The shEGFP(+) cells except ctxAB(++/2) cells comprised a cluster distinct from GFP(-) cells, supporting the notion that these cells have already undergone LTP-IE probably while the animal visited context A or B (potentiation of ratio, t = -5.985, p < 0.001; Baseline ratio, t = 7.899, p < 0.001, *k*-means clustering and independent t-test; Fig. S6E). The ctxAB(++/2) cells (n = 7) were not different from GFP(-) cells (potentiation of EPSP/EPSC ratio, t = -0.171, p = 0.868; baseline ratio, t = 0.154, p = 0.881, independent t-test; Fig. S6E).

#### *Summary for the ensemble dynamics in the CA3 network*

Based on our CA3 network model (see *Materials and Methods*), memories for two temporally separate events can be associated in the CA3 model network via the following mechanism:

1. *CA3-PCs primed by* $\boldsymbol{P_1}$ *are readily fired by PP input alone during the encoding phase of* $\boldsymbol{P_2}$: We assumed that a fraction of $\mathbf{M_1}$ PCs receives strong MF inputs during encoding phase of $\boldsymbol{P_1}$, and undergo LTP-IE, resulting in enhanced E-S coupling of subsequently incoming PP inputs. These primed PCs of $\mathbf{M_1}$ subset (denoted as $\mathbf{M_1}$***** PCs) are readily fired by PP inputs alone of the next pattern $\boldsymbol{P}_2$.
2. *Upon the arrival of* $\boldsymbol{P}_2$ *PP-LTP is induced not only in* $\mathbf{M_2}$ *PCs but also in* $\boldsymbol{M_1}$******* *PCs*: Sparse and decorrelated MF inputs of $\boldsymbol{P}_2$ create an $\mathbf{M_2}$ subset orthogonal to the $\mathbf{M_1}$ subset. The orthogonality of $\mathbf{M_1}$ and $\mathbf{M_2}$ subsets makes hetero-association of the two subsets unlikely when the two episodes occur in a temporally discontiguous sequence. However, PP inputs of $\boldsymbol{P_2}$ to the $\mathbf{M_1}$***** PCs readily elicit postsynaptic APs, which facilitate LTP induction at PP synapses carrying $\boldsymbol{P}_2$. Therefore, the PP inputs of $\boldsymbol{P_2}$ induce LTP not only in $\mathbf{M_2}$ but also in $\mathbf{M_1}$***** PCs. Furthermore, the primed state is reversed (depotentiation) by burst firing elicited by PP inputs as shown in Fig. 4. Thus, the $\mathbf{M_1}$***** PCs do not fire any longer in response to PP inputs in response to subsequent patterns.

3. *Recurrent connections are strengthened between PCs in **$M_2$** and **$M_1$****: ***P***$_2$ elicits firing both of **$M_2$** PCs and **$M_1$*** PCs, and thus recurrent connections are strengthened not only between **$M_2$** PCs but also between **$M_2$** and **$M_1$*** PCs. Therefore, the **$M_1$*** PCs play a key role in association between **$M_1$** and **$M_2$**, the two ensembles which encode ***P***$_1$ and ***P***$_2$, respectively.

### Supplementary Figure Legends

*Fig. S1*

**Fig. S1 Intracellular signaling cascade that mediates MF conditioning-induced downregulation of Kv1.2 in CA3-PCs based on our previous studies (Hyun et al., 2013, 2015; Eom et al. 2019)**. (1) High frequency MF input elicits back-propagating APs (bAPs), which induce dendritic $Ca^{2+}$ signaling through L-type voltage-dependent $Ca^{2+}$ channels (VDCC). (2) The dendritic $Ca^{2+}$ signaling activates protein tyrosine kinase (PTK), which induces endocytosis of Kv1.2 at distal apical dendrites. This downregulation of Kv1.2 underlies LTP-IE. (3) The dendritic Ca2+ signaling activates not only PTK but also receptor protein tyrosine phosphatase alpha (RPTPα), which antagonizes the PTK-dependent downregulation of Kv1.2. Thus, dendritic $Ca^{2+}$ signaling alone caused by bAPs upon somatic current injection induces LTP-IE only in a narrow range of $[Ca^{2+}]i$ (c.a. 340-380 nM, Eom et al., 2019). (4) High frequency MF input, however, induces intracellular $Zn^{2+}$ signaling together with bAP-mediated $Ca^{2+}$-signaling in postsynaptic CA3-PCs. The Zn2+ signaling inhibits RPTPα, and thus disinhibits the action of PTK leading to facilitation of LTP-IE.

*Fig. S2*

**Fig. S2**. **Test for stability of whole-cell patch recordings in the presence of 5 mM GSH in the pipette solution. Aa.** Time course for baseline-normalized input conductance ($G_{in}$) of CA3-PCs. Inset, Representative traces for measuring $G_{in}$ at the time indicated by arrowheads (black, 5 min; red, 45 min). **Ab.** Summary for $G_{in}$ of CA3-PCs. Reduced glutathione (GSH) in the patch pipette solution did not change $G_{in}$ of CA3-PCs over time (3.72 ± 0.42 nS to 3.87 ± 0.35 nS, n = 5, t = 0.223 p = 0.834, paired t-test). **Ba.** Baseline-normalized PP-EPSP amplitude. *Inset*, Representative traces for PP-EPSPs at the time indicated by arrowhead (black, 5 min; red, 45 min). **Bb**. Mean values for PP-EPSP amplitude. GSH in patch pipette solution did not change the PP-EPSP amplitude over time (0.66 ± 0.06 mV to 0.57 ± 0.11 mV, n = 5, t = 0.040, p = 0.970, paired t-test). Comparison of these $G_{in}$ and PP-EPSP amplitudes with those measured without GSH (reproduced from ref. 3) revealed that neither the baseline $G_{in}$ nor PP-EPSPs was altered by intracellular GSH ($G_{in}$ with GSH, 4.63 ± 1.47 nS, n = 14; $G_{in}$ without GSH, 3.75 ± 0.51 nS, n = 9; t = -1.722, p = 0.100, independent t-test; PP-EPSP with GSH, 0.56 ± 0.04 mV, n = 7; PP-EPSP without GSH, 0.66 ± 0.05 nS, n = 9; t = 1.512, p = 0.162 independent t-test). **C**. The latency of the first action potential (1st AP latency, *Ca-Cb*) and AP threshold (*Cc*) measured from voltage responses to a ramp current injection (250 pA/s) were not different between at 5 min and at 45 min of WCR. **Ca.** Representative membrane voltage responses to somatic ramp current injection (250 pA/s, black, at 5 min; red, at 45 min). **Cb.** The 1st AP latency (687.4 ± 78.73 ms to 662.4 ± 99.6 nS, n = 5, t = 0.362, p = 0.736, paired t-test). **Cc.** The AP threshold (-43.67 mV to -42.28 mV, n = 5, t = -1.16, p = 0.311, paired t-test).

*Fig. S3*

**Fig. S3**, **The somatic or MF conditioning lowers the threshold for homosynaptic induction of LTP of PP-EPSCs (PP-LTP) in CA3-PCs**. WCRs of CA3-PCs were done using GSH-free patch pipette solution. **Aa** and **Ba**. Baseline-normalized amplitudes of PP-EPSPs (evoked every 10 s) were monitored before and after high-frequency stimulation of PP-CA3 synapses (PP-HFS, red arrow) with weak (*Aa*) or strong (*Ba*) stimulation intensity. PP-HFS is comprised of 10 bursts every 10 s, and each burst of 20 stimuli at 100 Hz. The stimulation intensity was quantified by the baseline PP-EPSC amplitude (weak, < 12.55 pA; strong, > 12.55 pA). Upper red traces show voltage responses to PP-HFS. A single burst response was annotated as a gray box, and shown in the expanded time scale on the right. Note that weak PP-HFS induced neither APs (*Aa*) nor PP-LTP, whereas somatic APs were evoked during strong PP-HFS (*Ba*). EPSP amplitudes of non-failure events were normalized to the baseline value in the same cell. Recording times for the representative EPSC and EPSP traces (insets) are denoted in the main graph by the numbers (black, control; red, 30 min after PP-HFS). **Ab** and **Bb**. Mean amplitudes of PP-EPSC and PP-EPSP before and 30 min after PP-HFS in cases of PP-LTP induction failure (*Ab*), and in cases when LTP was induced (*Bb*). **C**. Plot of potentiation ratio of PP-EPSCs induced by PP-HFS as a function of the baseline EPSC amplitude, which is a metric of PP stimulation intensity. PP-LTP was induced in naïve CA3-PCs (gray symbols), and in CA3PCs that underwent MF (red) or somatic (blue) conditioning delivered before PP-HFS. A logistic function (baseline: 108.14 ± 16.8%, max: 201.75 ± 29.6%, slope: 0.35578 ± 0.734, $EPSC_{50}$ = 13.077 ± 0.635 pA) was fitted to the LTP level of naïve CA3-PCs. **Da**. Voltage responses to weak PP-HFS in the CA3-PCs which underwent MF (upper, red) or somatic conditioning (lower, blue). The response to a single burst stimulation annotated as gray box is shown in the expanded time, in the right of each voltage responses. **Db**. PP-EPSC amplitude changes caused by MF or somatic conditioning (blue arrowhead) and subsequent PP-HFS (indicated by gray arrow at 0 min). Note that PP-LTP is readily induced by weak PP-HFS once the CA3-PC underwent MF (red symbols) or somatic (blue symbols) conditioning. **Dc**. Representative traces of PP-EPSCs (left) and PP-EPSPs (right) recorded at the times as indicated by the numbers in *Db* (black, baseline; blue, after MF (upper) or somatic (lower) conditioning; red, after PP-HFS). Note that MF or somatic conditioning enhanced PP-EPSPs but not PP-EPSCs, while both were enhanced by PP-HFS. **Ea-Eb**. Mean amplitudes of PP-EPSCs measured at the baseline, after MF (*Ea*) or somatic (*Eb*) conditioning, and after PP-HFS (conditioning: $F_{(1,19)}$ = 0.065, p = 0.802, time: $F_{(1,19)}$ = 74.965, p < 0.001, time × conditioning: $F_{(1,19)}$ = 0.012, p = 0.915; GLM and simple effect analysis). **Ec-d**. Mean amplitudes of PP-EPSPs measured at the baseline, after MF (*Ec*) or somatic conditioning (*Ed*), and after PP-HFS (conditioning: $F_{(1,19)}$ = 3.856, p = 0.064, time: $F_{(1,19)}$ = 11.858, p < 0.001, time × conditioning: $F_{(1,19)}$ = 0.886, p = 0.358; GLM and simple effect analysis).

*Fig. S4*

**Fig. S4 Analysis of Fluorescence intensity of shEGFP(+) CA3-PCs. A**. Schematic timeline of the experimental procedure. *Upper*: Mice visited context A for 5 min and then returned to homecage for 85 min, and sacrificed for *ex vivo* study (denoted as ctxA). *Lower*: After visiting context A for 5 min and being kept in homecage for 25 min, mice visited context B or C for 5 min, and then were kept in homecage for 55 min before ex vivo study (denoted as ctxAB). **B**. Maximum intensity projection of Z-stack confocal images ($Z_{max}$ image) of hippocampal CA3 region obtained from ctxA mice (Ba) and ctxAB mice (Bc). The right graph of each image shows the fluorescence intensity profiles of boxed GFP(+) CA3-PCs as a function of depth from a surface the slices. The same color codes are used for each box and corresponding profile. Fluorescence signals above a threshold of 5× the SD of baseline fluctuation were regarded as a fluorescence signal of GFP(+) CA3-PCs. **C**. Plot of maximum fluorescence intensity ($F_{max}$) of individual GFP(+) CA3-PCs as a function of the depth at which $F_{max}$ of each cell was measured. Distribution of GFP fluorescence measured in GFP(+) CA3-PCs from ctxAB or ctxAC mice was bimodal, in contrast to those of ctxA mice, and thus CA3-PCs from ctxAB or ctxAC mice were clustered into two groups: cells displaying low or high GFP fluorescence. **Cb**. Mean fluorescence of GFP(+) CA3-PCs measured in different cell groups. There was no significant difference in the fluorescence intensity between high fluorescence ctx AB and ctx AC cells. **D.** Histogram of fluorescence intensity of shEGFP(+) CA3-PCs from either ctxA (*Da*) or ctxAB (*Db*) mice. In each plot, mono- and bi-lognormal fits are shown. Mono- and bi-lognormal fits were better for ctxA and ctxAB, respectively. The best fit parameters of log-normal function [$y = A*\exp[-(\ln(x/m)/s)^2]$] for ctxA: A = 11.6, m = 37.6, s = 1.24. Those for ctxAB, $A_1$ = 7.41, $m_1$ = 55.87, $s_1$ = 1.106, $A_2$ = 1.497, $m_2$ = 521.6, s2 = 0.29.

*Fig. S5*

**Aa-b.** Representative traces for single AP evoked by short current injection and mean AP onset time before and after somatic conditioning (condition, $F_{(5,27)}$ = 2.465, p = 0.058; time, $F_{(1,27)}$ = 8.011, p = 0.009; condition x time: $F_{(5,27)}$ = 7.633, p < 0.001; GLM and simple effect analysis). **Ba-b.** Representative traces for first AP latency by ramp current injection and summary for change of first AP latency (condition, $F_{(5,27)}$ = 3.044, p = 0.026; time, $F_{(1,27)}$ = 11.951, p = 0.002; condition x time, $F_{(5,27)}$ = 4.570, p = 0.004). AP onset time compared to GFP(-) cells, ctxA60(+), $p < 0.001$; ctxA90(+), $p < 0.001$; ctxAB(+), $p < 0.001$. For first AP latency, ctxA60(+), $p < 0.001$; ctxA90(+), $p < 0.001$; ctxAB(+), $p < 0.001$.

*Fig. S6*

**Fig. S6**, Depotentiation of PP-EPSPs in twice-activated ensemble cells. **Aa.** Baseline EPSP/EPSC ratio as a function of G/R fluorescence ratio. **Ab.** Potentiation of EPSP/EPSC ratio (ordinate) is closely correlated with baseline $G_{in}$ (abscissa). Color codes for GFP(-), gray; ctxA60(+), blue; ctxA90(+), green; ctxAB(+), orange; ctxAB(++/1), violet; ctxAB(++/2), red. **Ba-b.** Representative traces for PP-EPSCs (*a*) and EPSPs (b) before (gray) and after (colored) somatic conditioning. The same color codes as in *A*. **Bc.** Time courses of PP-EPSP/EPSC ratio changes caused by somatic conditioning (large blue arrowhead at 0 min). *Inset*, AP train induced by the somatic conditioning. Representative traces shown in a-b were recorded at time points indicated by small arrowheads (black, baseline; red, 25 min). The same color codes as in *A* (condition: $F_{(5,33)} = 6.657$, $p < 0.001$; time: $F_{(1,33)} = 20.375$, $p < 0.001$; condition x time: $F_{(5,33)} = 8.813$, $p < 0.001$; GLM and simple effect analysis). **C.** Summary for PP-EPSC (*a*) and EPSP (*b*) amplitudes before (gray) and after (red) somatic conditioning. For EPSC, conditioning, $F_{(5,33)} = 0.406$, $p = 0.841$; time, $F_{(1,33)} = 0.130$, $p = 0.720$; condition × time, $F_{(5,33)} = 0.891$, $p = 0.498$. For EPSP, conditioning, $F_{(5,33)} = 2.385$, $p = 0.059$; time, $F_{(1,33)} = 21.676$, $p < 0.001$; condition × time, $F_{(5,33)} = 13.016$, $p < 0.001$; RM-ANOVA. For pre- vs post-SC EPSPs (*Cb*), ctxA60(+), $p = 0.33$; ctxA90(+), $p = 0.160$; ctxAB(+), $p = 0.811$; ctxAB(++/1), $p = 0.368$; ctxAB(++/2), $p < 0.001$; GFP(-), $p < 0.001$. **D.** PP-EPSP amplitude as a function of PP-EPSC of each CA3-PC. The EPSP/EPSC ratio of CA3-PCs were categorized into two groups: high and low slope groups ($F_{(6,39)} = 17.181$, $p < 0.001$, 1-way ANOVA and Tukey HSD). Note that the post-SC EPSC/EPSC ratio of GFP(-) cells belonged to the high slope group. **E**. Somatic conditioning-induced potentiation of EPSP/EPSC ratio as a function of its baseline value. CA3-PCs were classified using k-mean analysis. There was a significant difference between ctxAB(GFP++/1) and ctxAB(GFP++/2) groups. The same color codes as in *A*.

*Fig. S7.*

**Fig. S7, A**. Connectivity map between PCs and INs (blue, not connected; light blue, connected). PCs are indexed from 1 to 1000, and INs are from 1001 to 1200. The baseline synaptic conductance between PCs ($J_{ee}$) was set to a random value between 0 and 0.35 nS. The synaptic conductance values for PCs-to-INs ($J_{ie}$), INs-to-PCs ($J_{ei}$), INs-to-INs ($J_{ii}$) were set to 0.4 nS. c, connection probability; J, baseline synaptic weight; e, excitatory; i, inhibitory; $J_{YX}$, synaptic conductance from X to Y. **Ba-Bb**, STDP rule at A/C and PP synapses. The LTP ratio (L), was modelled as, $L(\Delta t) = 2 \exp(-|\Delta t|/40)$ for A/C synapses. For PP synapses, $L(\Delta t) = 2$

exp(-Δt/30) if $\Delta t \geq 0$, and -0.5 exp(Δt/15) if $\Delta t < 0$. **C**, To emulate a distal dendritic spike elicited by PP inputs to a primed PC (Hyun et al., 2015), the inward current pulse was injected upon arrival of PP inputs in a primed PC (Jahnke et al., 2015). **D**, Distributions of the number of MF and PP inputs to each CA3-PC per a pattern. Left, the number of MF inputs to a CA3-PC in an active **M** subset was set to a random number which follows log-normal distribution with $\mu = 1$ and $\sigma = 0.6$. The MF input numbers were bounded by [3, 15]. Right, Distributions for the number of PP inputs during recall and encoding phases are compared (red and blue bars, respectively). The PP input numbers were also set by random values of lognormal distribution with $\mu = 1.8$ and $\sigma = 0.3$ for encoding phase [number of inputs, 6.33 $\pm$ 3.77 (mean $\pm$ variance)], and by those with $\mu = 1.2$ and $\sigma = 0.3$ for recall phase [number of inputs, 3.47 $\pm$ 1.14]. The number of PP inputs was bounded by [3, 20] for encoding, and by [0, 10] for recall. **E**, The cue input incoming via PP that triggers retrieval of $\mathbf{M_1}$. The PP inputs are light-blue colored if the synaptic weights have been strengthened during the encoding phase, and magenta otherwise.

Fig. S1

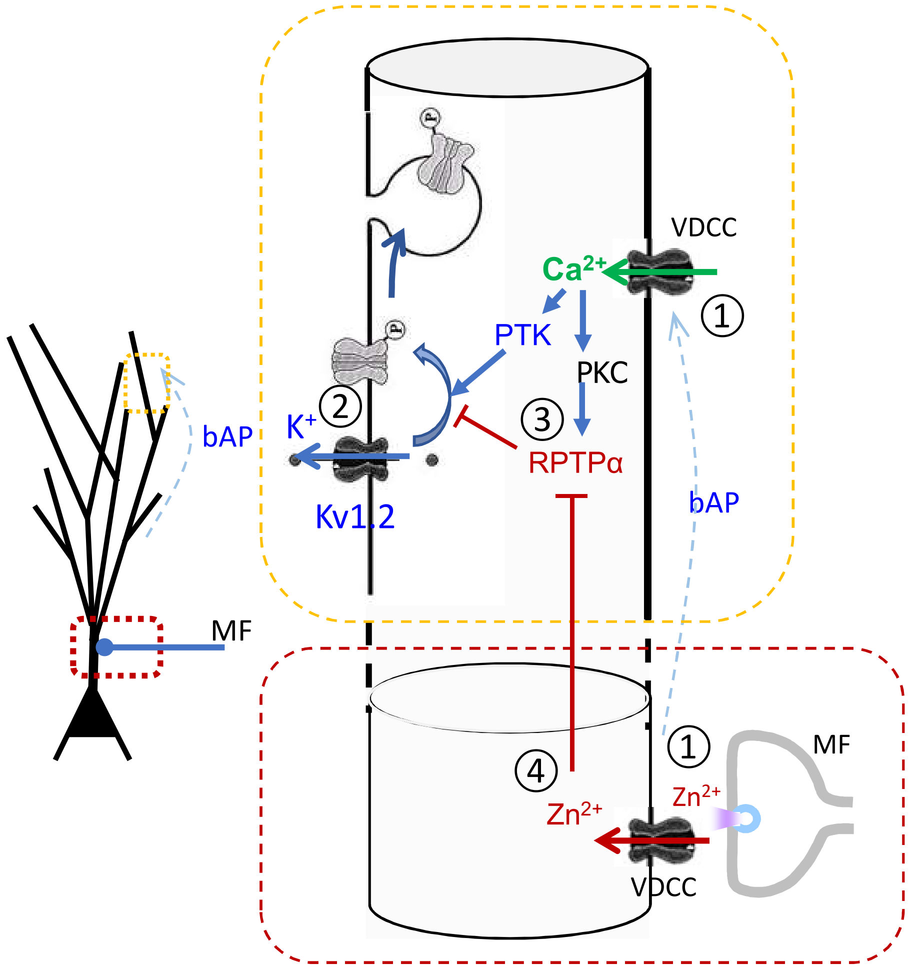

Fig. S2

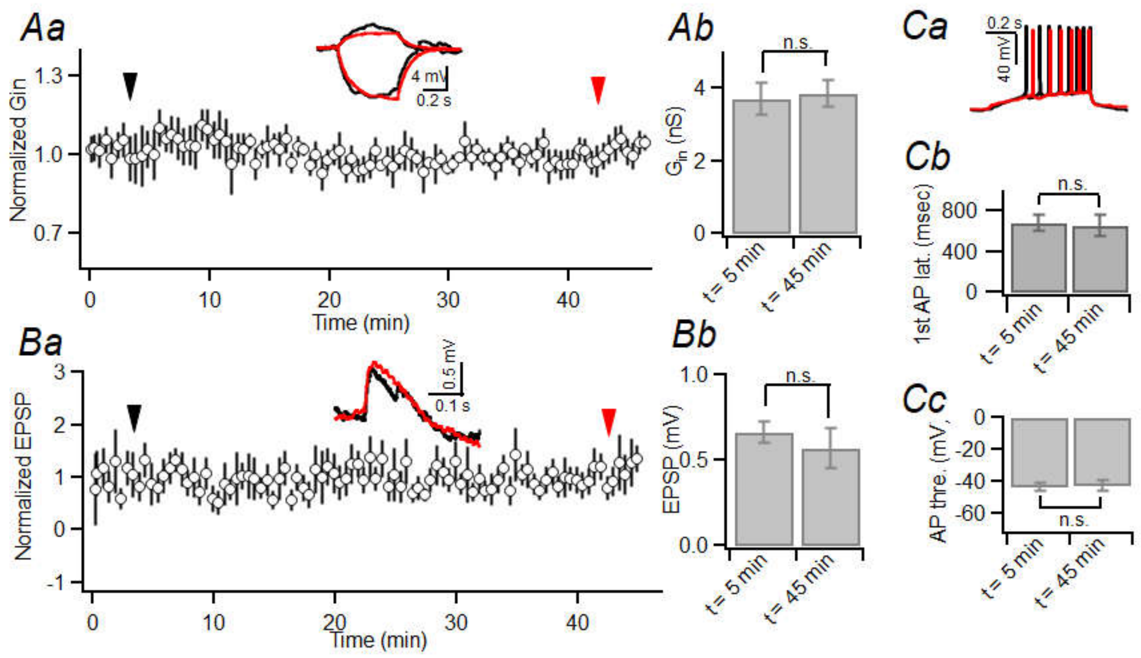

Aa
Ab
Ca
Cb
Cc
Ba
Bb
Normalized Gin
Time (min)
4 mV
0.2 s
n.s.
Gin (nS)
t= 5 min
t= 45 min
0.2 s
40 mV
1st AP lat. (msec)
Normalized EPSP
0.5 mV
0.1 s
EPSP (mV)
AP thre. (mV)

Fig. S3

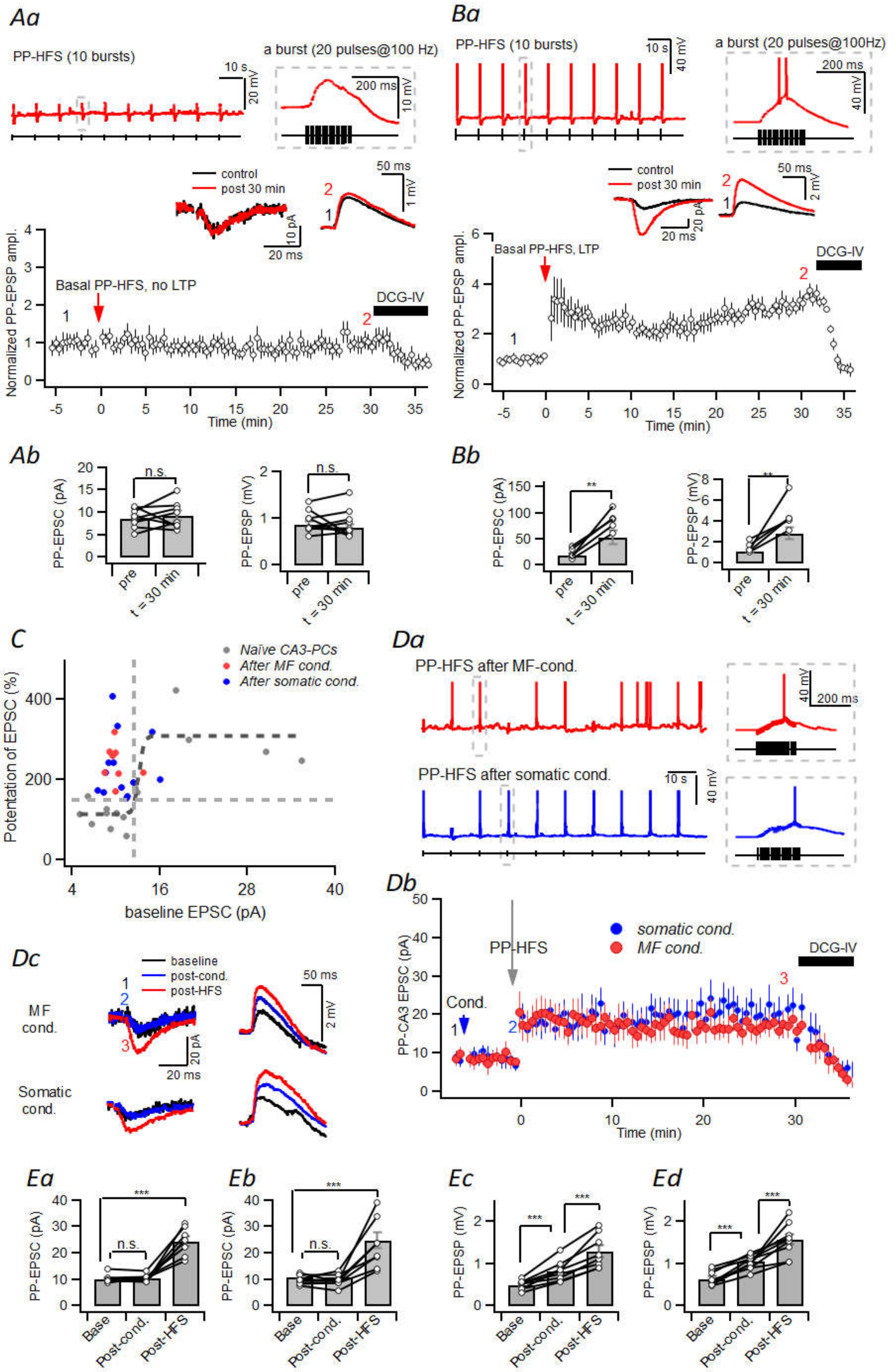

Aa
PP-HFS (10 bursts)
a burst (20 pulses@100 Hz)
control
post 30 min
Basal PP-HFS, no LTP
DCG-IV
Normalized PP-EPSP ampl.
Time (min)
Ba
PP-HFS (10 bursts)
a burst (20 pulses@100Hz)
Basal PP-HFS, LTP
Ab
PP-EPSC (pA)
PP-EPSP (mV)
n.s.
pre
t = 30 min
Bb
C
Naïve CA3-PCs
After MF cond.
After somatic cond.
Potentiation of EPSC (%)
baseline EPSC (pA)
Da
PP-HFS after MF-cond.
PP-HFS after somatic cond.
Db
PP-CA3 EPSC (pA)
PP-HFS
somatic cond.
MF cond.
Cond.
Dc
baseline
post-cond.
post-HFS
MF cond.
Somatic cond.
Ea
Eb
Ec
Ed
Base
Post-cond.
Post-HFS

Fig. S4

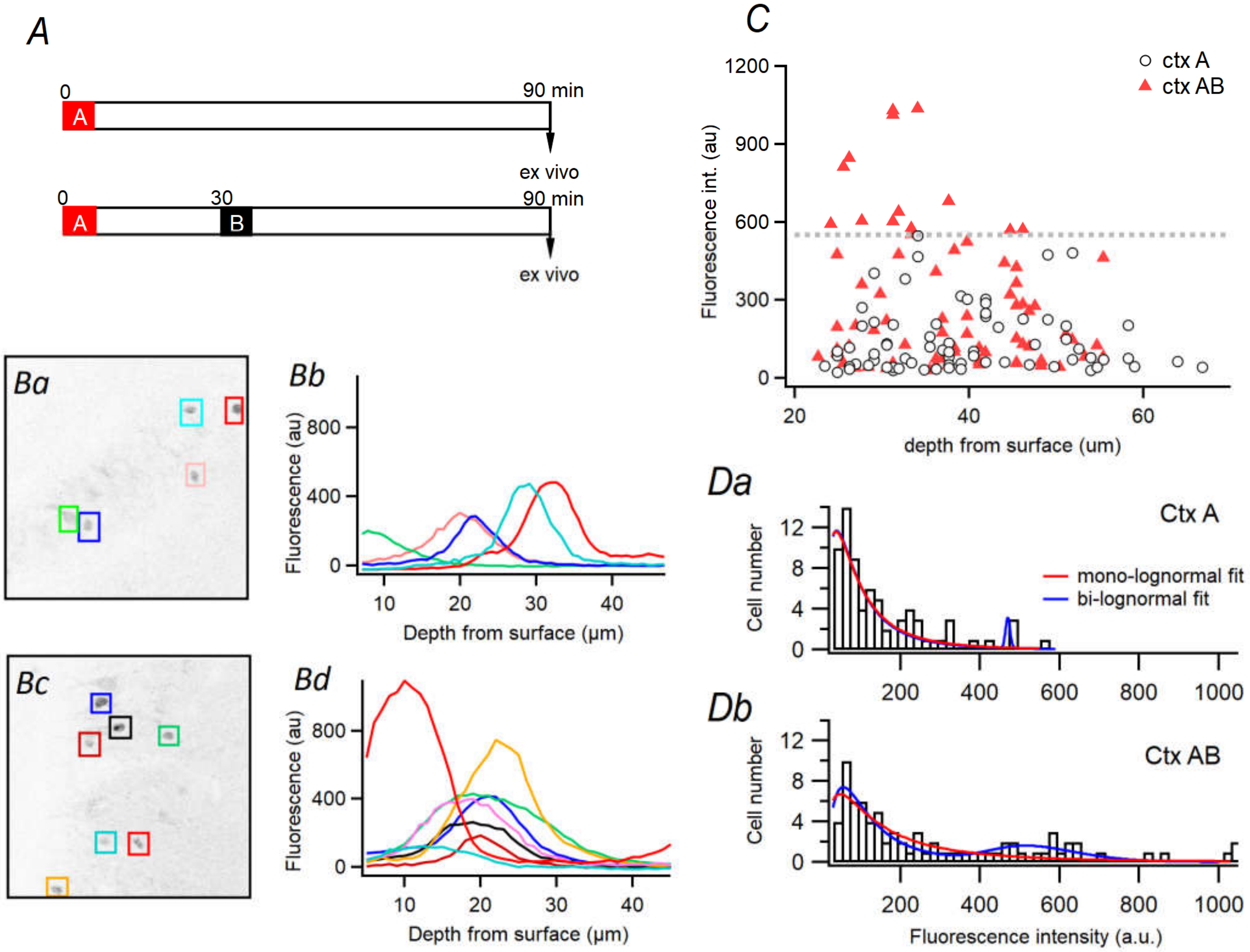
A
0
90 min
A
ex vivo
0
30
90 min
A
B
ex vivo
Ba
Bb
Fluorescence (au)
Depth from surface (µm)
Bc
Bd
Fluorescence (au)
Depth from surface (µm)
C
ctx A
ctx AB
Fluorescence int. (au)
depth from surface (um)
Da
Ctx A
mono-lognormal fit
bi-lognormal fit
Cell number
Db
Ctx AB
Cell number
Fluorescence intensity (a.u.)

Fig. S5

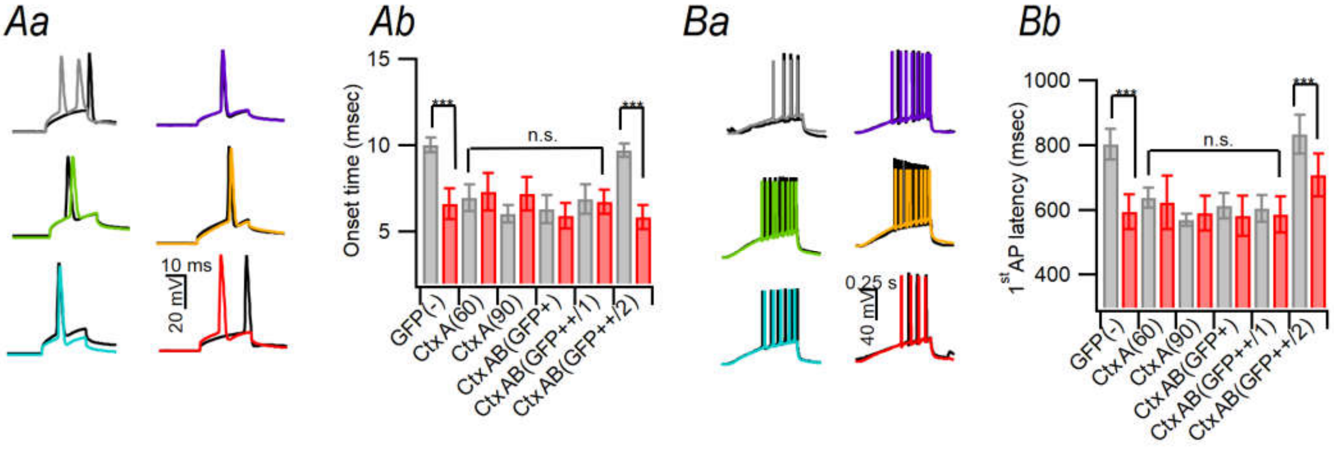

Aa
Ab
Ba
Bb
10 ms
20 mV
15
10
5
Onset time (msec)
***
n.s.
GFP(-)
CtxA(60)
CtxA(90)
CtxAB(GFP+)
CtxAB(GFP++/1)
CtxAB(GFP++/2)
0.25 s
40 mV
1000
800
600
400
1st AP latency (msec)

Fig. S6

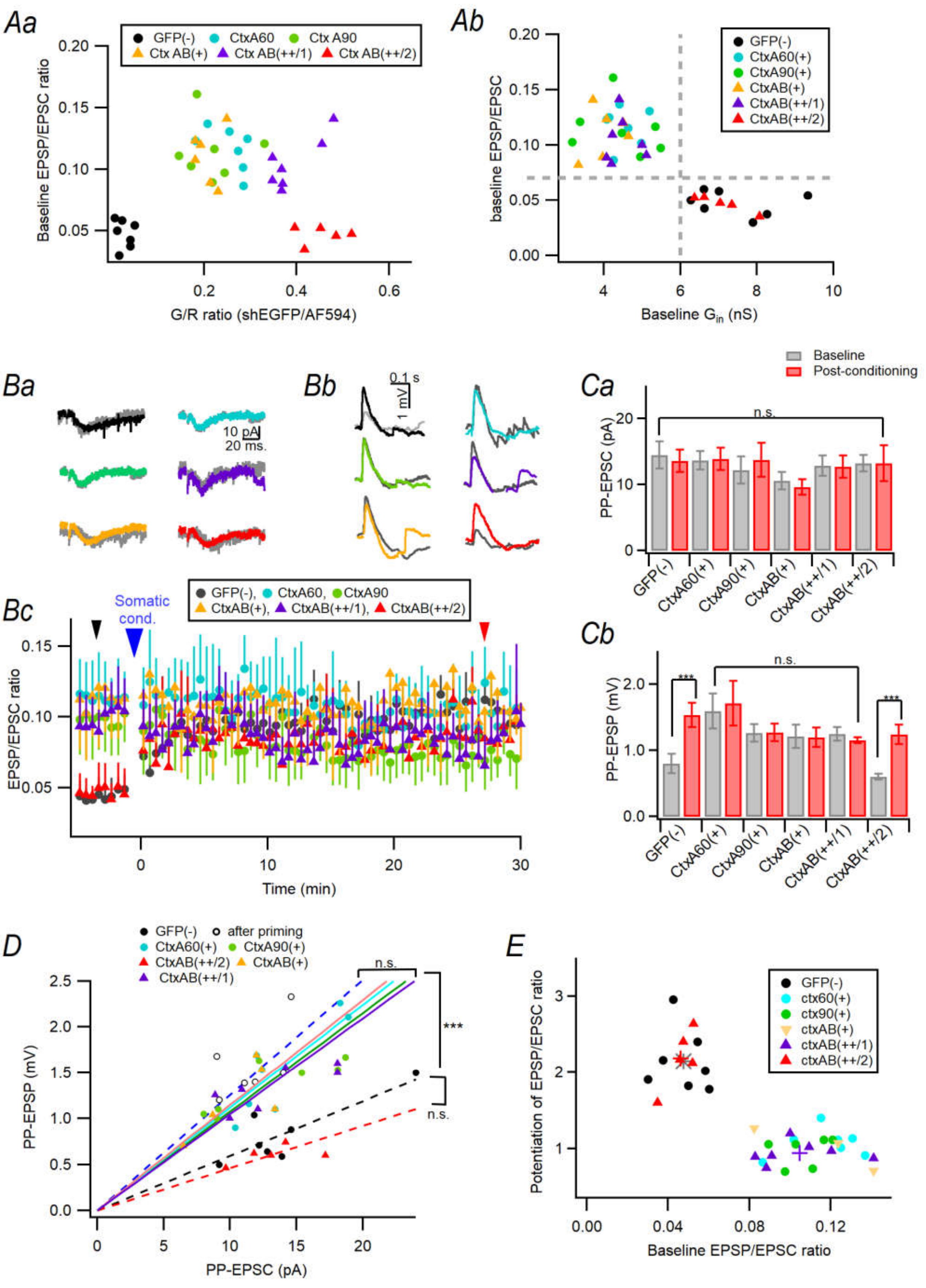

Aa
GFP(-) CtxA60 Ctx A90
Ctx AB(+) Ctx AB(++/1) Ctx AB(++/2)
Baseline EPSP/EPSC ratio
G/R ratio (shEGFP/AF594)
Ab
GFP(-)
CtxA60(+)
CtxA90(+)
CtxAB(+)
CtxAB(++/1)
CtxAB(++/2)
baseline EPSP/EPSC
Baseline $G_{in}$ (nS)
Ba
10 pA
20 ms.
Bb
0.1 s
1 mV
Ca
Baseline
Post-conditioning
n.s.
PP-EPSC (pA)
GFP(-) CtxA60(+) CtxA90(+) CtxAB(+) CtxAB(++/1) CtxAB(++/2)
Bc
Somatic cond.
GFP(-), CtxA60, CtxA90
CtxAB(+), CtxAB(++/1), CtxAB(++/2)
EPSP/EPSC ratio
Time (min)
Cb
***
n.s.
***
PP-EPSP (mV)
GFP(-) CtxA60(+) CtxA90(+) CtxAB(+) CtxAB(++/1) CtxAB(++/2)
D
GFP(-) after priming
CtxA60(+) CtxA90(+)
CtxAB(++/2) CtxAB(+)
CtxAB(++/1)
n.s.
***
n.s.
PP-EPSP (mV)
PP-EPSC (pA)
E
GFP(-)
ctx60(+)
ctx90(+)
ctxAB(+)
ctxAB(++/1)
ctxAB(++/2)
Potentiation of EPSP/EPSC ratio
Baseline EPSP/EPSC ratio

Fig. S7

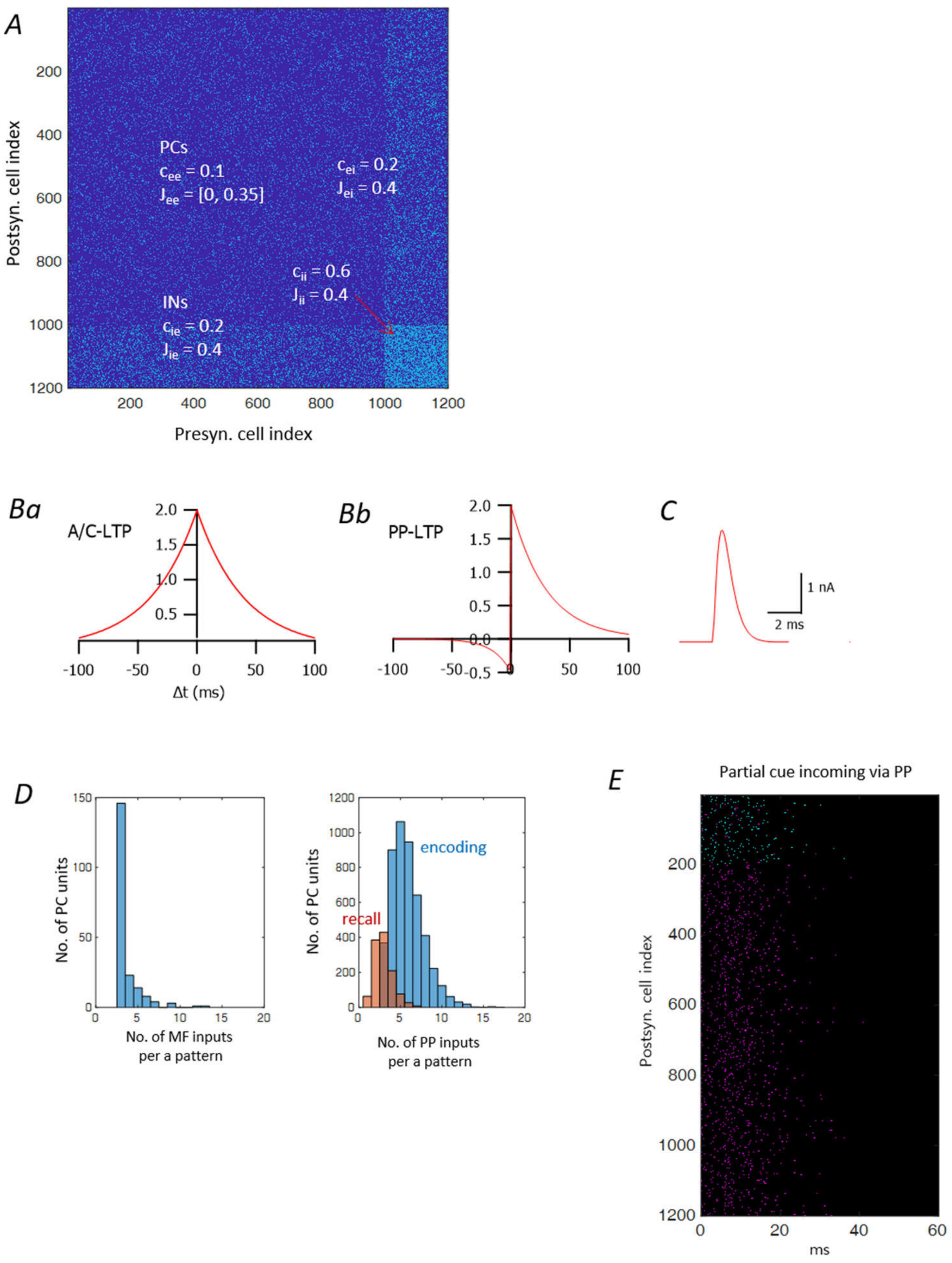
A
PCs
$c_{ee}$ = 0.1
$J_{ee}$ = [0, 0.35]
$c_{ei}$ = 0.2
$J_{ei}$ = 0.4
$c_{ii}$ = 0.6
$J_{ii}$ = 0.4
INs
$c_{ie}$ = 0.2
$J_{ie}$ = 0.4
Postsyn. cell index
Presyn. cell index
Ba
A/C-LTP
Δt (ms)
Bb
PP-LTP
C
1 nA
2 ms
D
No. of PC units
No. of MF inputs per a pattern
encoding
recall
No. of PP inputs per a pattern
E
Partial cue incoming via PP
Postsyn. cell index
ms
